# Patient-derived recombinant autoantibodies reveal the subcellular neuroglial distribution and heterogeneous interactome of leucine-rich glioma-inactivated 1

**DOI:** 10.1101/2022.03.29.486232

**Authors:** Jorge Ramirez-Franco, Kévin Debreux, Johanna Extremet, Yves Maulet, Maya Belghazi, Claude Villard, Marion Sangiardi, Fahamoe Youssouf, Lara El Far, Christian Lévêque, Claire Debarnot, Pascale Marchot, Sofija Paneva, Dominique Debanne, Michael Russier, Michael Seagar, Sarosh R Irani, Oussama El Far

**Affiliations:** UMR 1072, INSERM, Unité de Neurobiologie des canaux Ioniques et de la Synapse, (UNIS) Aix-Marseille Université, 13015 Marseille, France; Aix-Marseille Univ, CNRS, Institute of Neurophysiopathology (INP), PINT, PFNT, 13385 Marseille cedex 5, France; Lab ’Architecture et Fonction des Macromolécules Biologiques (AFMB)’, CNRS, Aix-Marseille Université, 13288 Marseille cedex 09, France; Oxford Autoimmune Neurology Group, Nuffield Department of Clinical Neurosciences, John Radcliffe Hospital, University of Oxford, Oxford, UK; Department of Neurology, Oxford University Hospitals, Oxford, UK

**Keywords:** Leucin-rich Glioma inactivated 1 (LGI1) - Auto-antibodies - Limbic Encephalitis - Autosomal-dominant lateral temporal lobe epilepsy (ADLTE) - Kv1 channels - ADAM

## Abstract

Autoantibodies against leucine-rich glioma-inactivated 1 (LGI1) occur in patients with encephalitis who present with frequent focal seizures and a pattern of amnesia consistent with focal hippocampal damage. To investigate whether the cellular and subcellular distribution of LGI1 may explain the localisation of these features, and gain broader insights into LGI1 neurobiology, we analysed the detailed localisation of LGI1, and the diversity of its protein interactome, in mouse brains using recombinant monoclonal LGI1-antibodies derived from encephalitis patients. Combined immunofluorescence and mass spectrometry analyses showed that LGI1 is enriched in excitatory and inhibitory synaptic contact sites, most densely within CA3 regions of the hippocampus. LGI1 is secreted in both neuronal somatodendritic and axonal compartments, and occurs in oligodendrocytic, neuro- oligodendrocytic and astro-microglial protein complexes. Proteomic data support the hypothesis that destabilization of Kv1 / MAGUK complexes by autoantibodies could promote excitability, but did not reveal LGI1 complexes with postsynaptic glutamate receptors. Our results extend our understanding of regional, cellular and subcellular LGI1 expression profiles and reveal novel LGI1-associated complexes, thus providing insights into the complex biology of LGI1 and its relationship to seizures and memory loss.

## Introduction

LGI1 (Leucine-rich Glioma Inactivated 1) is a secreted glycoprotein, predominantly expressed in the central nervous system. It is composed of an N-terminal leucine-rich repeat (LRR) and a C-terminal epitempin (EPTP) domains. Through the latter it interacts with the ectodomains of ADAM22, 23 and 11 which are catalytically-inactive members of the ADAMs family of metalloproteases [80] [19] [65] [56]. It has been suggested that LGI1 plays an essential role in the formation of a functional trans-synaptic complex linking presynaptic Kv1 channels to postsynaptic AMPA/NMDA receptors and PSD95 through interaction with ADAM 22/23 [19] [21] [46] [20].

LGI1 owes its name to its downregulated expression in malignant gliomas, its first involvement in human disease [8]. Point mutations in LGI1, the majority of which lead to an inhibition of LGI1 secretion [69] [52] [45], have been linked to inherited, autosomal-dominant lateral temporal lobe epilepsy (ADLTE) [51] [34] and neuronal hyperexcitability [83] [5] [87]. In ADLTE, the few secreted LGI1 mutants show impaired ADAM22 binding [12] [81], suggesting that hypofunction of the ADAM22-LGI1 interaction may account for the phenotype associated with these human LGI1 mutations [20]. In mice, LGI1 deletion results in lethal seizures accompanied by a reduction in the AMPA / NMDA receptor ratio and a decrease in AMPA receptor-mediated synaptic transmission [21] [22] [46]. However, whether these modifications account for the observed neuronal hyperexcitability is unclear [83] [5] [87]. We recently showed that downregulation of Kv1 channels in axonal initial segments (AIS) may also contribute to epileptogenesis in *Lgi1^-/-^* mice, through an increase in the intrinsic excitability of CA3 neurons [68], and suggested that the decrease in AMPA receptors may be a consequence of neuronal hyperexcitability. Furthermore, hypomyelination [71] and alterations in dendritic pruning and spine development [87] [75] have been described in *Lgi1^-/-^* mice. More recently, a novel missense LGI1 mutation was shown to impair oligodendrocyte differentiation and induce white matter abnormalities [74]. Collectively, these findings suggest that LGI1 is differentially involved at distinct developmental stages, in a cell and context-specific manner [10].

The link between LGI1 and Kv1 channels has been supported by immunoprecipitation studies using anti Kv1.2 antibodies [53] or human LGI1-autoantibodies [30] [41] [48]. Upon passive transfer to rodents, LGI1 antibodies modulate both Kv1 function and expression and AMPAR function, akin to genetic paradigms [58] [63]. Furthermore, when administrated in animal models, these antibodies reproduce key cognitive features of the patients from which they were derived – namely a marked, and partially reversible amnesia related to the CA3-predominant, hippocampal damage observed in patients with LGI1-antibody encephalitis [49] [50] [58] [76].

Throughout these studies, examining the role of native LGI1 has been challenging, partly because available commercial and custom made LGI1-antibodies are of poor quality [25] [4] [5] and rarely validated on knockout tissues. Thus, they represent unreliable tools to define the native LGI1 interactome or its brain distribution. The search for native partners of LGI1 has been addressed using recombinant, tagged LGI1 either by adding it to brain extracts [39] or by generating knock-in mice [21]. These approaches showed an association of LGI1 with synapse-related protein complexes. Also, aspects of LGI1’s neurobiology have been partially addressed in dissociated hippocampal cultures [27] but not in structured tissues such as organotypic slices. Furthermore, different methodological approaches have resulted in contradictory results regarding LGI1’s expression pattern. For example, using GFP-expressing cells under the control of a cis-regulatory element of LGI1 in mice, a widespread distribution of LGI1 from embryonic to late postnatal stages has been determined [25] [72]. Whereas, analysis of mRNA distribution in adult mice brains, and other approaches, showed that LGI1 is highly enriched in the hippocampal circuitry with strong expression in CA3 and the dentate gyrus (DG) [26] [64] [7].

The most accurate and specific staining of native LGI1 to date has been achieved using patient-derived LGI1-antibodies, which are directly pathogenic *in vivo* [41] [54] [58]. However, these studies mainly addressed regional LGI1 distribution and did not focus on potential enrichment in subcellular domains and pre- or post-synaptic processes.

Recently, we isolated and characterized several monoclonal antibodies from two patients with LGI1- antibody encephalitis, which targeted either its native LRR or EPTP domains [63]. In the present study, we used these patient-derived domain-specific monoclonal-antibodies along with WT and *Lgi1^-/-^* animal tissues, as well as exogenously expressed fluorescent LGI1, to characterize, in the murine hippocampus, the detailed subcellular distribution and secretion pattern of LGI1. Furthermore, by immunoprecipitation and mass spectrometry from WT and *Lgi1^-/-^* brain extracts, we identified a series of distinct molecular complexes which constitute the native LGI1 interactome. Our findings reconcile a large number of contradictory findings concerning the functional consequences of LGI1 depletion and offer insights into the localisation and function of this enigmatic protein, whose dysfunction is increasingly implicated in human diseases.

## Materials and Methods

Methods are described below, briefly. Further details are available online as Electronic Supplementary Materials.

### Construction of plasmids for inducible LGI1 expression in mammalian-cells

A smaller version (pIND-IRES-EGFP) of the pINDUCER11 (miR-RUG) plasmid [47] was first generated and pIND-LGI1-IRES-EGFP and pIND-ΔIRES-Dendra2-LGI1 were constructed as described in the Electronic Supplementary Materials.

### LGI1 knock-out mice

Heterozygous *Lgi1*^+/−^ mice [7] were inter-crossed to generate *Lgi1^−/^*^−^ and *Lgi1^+/+^* littermates. All experiments were carried out according to the European and Institutional guidelines for the care and use of laboratory animals (Council Directive 86/609/EEC and French National Research Council) and were approved by the local animal health authority (# D 13 055 08, Préfecture des Bouches-du-Rhône, Marseille). This study does not involve experiments on live animals.

### Antibodies

A set of recombinant monoclonal IgG LGI1-antibodies consisting in three LRR-directed (herein referred to as LRR1-3; mAb01, 02 and 07 respectively in [63]) and two non-previously described EPTP-directed (referred to as EPTP1-2) was generated from two encephalitis patients, as previously described [63]. Other antibodies used in this study are described in Electronic Supplementary Materials.

### Surface Plasmon Resonance

Using a Biacore T200 and a CM5_sensor chip (Cytiva), purified His-tagged recombinant LGI1 [68] (about 1-2 fmoles) was captured on covalently coupled EPTP2. EPTP1, EPTP2, LRR1, or healthy control (HC) human IgGs were then injected at 1 μl/min for 5 min at different concentrations. Background signals were subtracted from flow cells in which irrelevant human IgGs were immobilized.

### Immunohistofluorescence of fixed brains

P14-P16 C57BL/6 wild-type mice or *Lgi1*^-/-^ littermates of either sex were used and prepared as described in the Electronic Supplementary Materials.

### Hippocampal slice cultures and transfection of hippocampal neurons

Hippocampal organotypic slices were prepared as previously described [61]. Details and the transfection procedure are described in the Electronic Supplementary Materials.

### Immunofluorescence of organotypic slices and image analysis

For the immunolabeling of organotypic slices, we adapted the protocol described in [24]. Images were analyzed with ImageJ (NIH). Details are described in the Electronic Supplementary Materials.

### Statistics

Statistical analysis was performed with Origin 8.0 and the specific test is indicated in the figure legends. One-way ANOVA followed by Bonferroni’s post-hoc test was used for multiple comparisons. Comparisons were considered significant when p-value<0.05. Figures were prepared for presentation using Adobe Illustrator.

### Immunoprecipitations

Wild type and *Lgi1^−/^*^−^ brains were extracted, snap-frozen in liquid nitrogen and kept at -80°C until use. Brains were homogenised in HB buffer (20 mM NaP pH 7.4, 30 mM NaCl containing phosphatase inhibitors (Pierce^TM^) and protease inhibitors (Complete, Roche Diagnostics GmbH)). Each brain was homogenised using 1 ml of HB and nuclei were eliminated by 900 x g centrifugation. Supernatants were solubilized in HB supplemented with 1% CHAPS at 5 mg/ml and subjected to 100.000 x g ultracentrifugation. The solubilised material from both wildtype and LGI1 knockout samples were precleared with rProtein A Sepharose Fast Flow (Cytiva) and immunoprecipitated in triplicates using the human autoimmune IgGs (LRR1; LRR3, EPTP1; EPTP2) (3-5 µg / immunoprecipitated sample).

Immunoprecipitated samples were washed 3 times with HB containing 0.5% CHAPS before denaturation. Samples were then analysed either by SDS-PAGE and Western blot or by mass spectrometry.

### Mass spectrometry analysis and data processing

Samples were analyzed by mass spectrometry using a hybrid Q-Orbitrap mass spectrometer as described in [61] (Q-Exactive, Thermo Fisher Scientific, United States) coupled to a nanoliquid chromatography (LC) Dionex RSLC Ultimate 3000 system (Thermo Fisher Scientific, United States). For details see the Electronic Supplementary Materials.

## Results

### Native LGI1 distribution in mouse brain

To characterize the expression pattern of native LGI1 in mice brain, we used three patient-derived LRR- directed antibodies of which two are competing (LRR1 & LRR2) [63]. Similar to mRNA distribution [26], immunofluorescence in coronal telencephalic slices revealed equivalent staining patterns with all three LRR-antibodies which was abrogated in *Lgi1^-/-^* sections (Fig. 1a-b). LGI1 expression intensities, displayed as a Look Up Table (LUT) for each antibody and normalized to hippocampal levels, showed LGI1 is most concentrated in the hippocampus and caudate-putamen complex, with far lower expression intensities in other cortical regions and basal ganglia / diencephalic regions (Fig. 1b-c). An intra- hippocampal, analysis showed ∼50% less LGI1 expression in ventral versus dorsal CA3 (Fig. 2a and b). Further analysis of sub-regional distribution of LGI1 in the dorsal hippocampus revealed its prominent expression in CA1, CA3 and DG but barely detectable levels in the Stratum Pyramidale (SP) and Granule Cell Layer (GCL) (Fig 2). Normalized immunoreactivity (Fig. 2b-e) shows a pattern of dorsal expression where CA3 = DG > CA1, and that the highest LGI1 density is observed at the level of the CA3 stratum radiatum (CA3_SR), where afferents from contralateral CA3 and recurrent collaterals of ipsilateral CA3 pyramidal neurons establish synapses with the distal apical dendrites of CA3 pyramidal neurons. Among the three main CA3 strata, LGI1 expression was lower in CA3 Stratum Lucidum (CA3_SL) where mossy fibres (MF) meet the proximal apical dendrites and thorny excrescences of CA3 pyramidal neurons. Furthermore, these differential regional expression levels were also observed when the punctate LGI1 cluster densities were studied (Fig. 2d, f), with significantly higher cluster densities in CA3_SR and DG ML versus CA1 subsectors, corroborating imaging studies of hippocampal damage in human patients with LGI-mediated limbic encephalitis [50] [49].

**Fig. 1.**
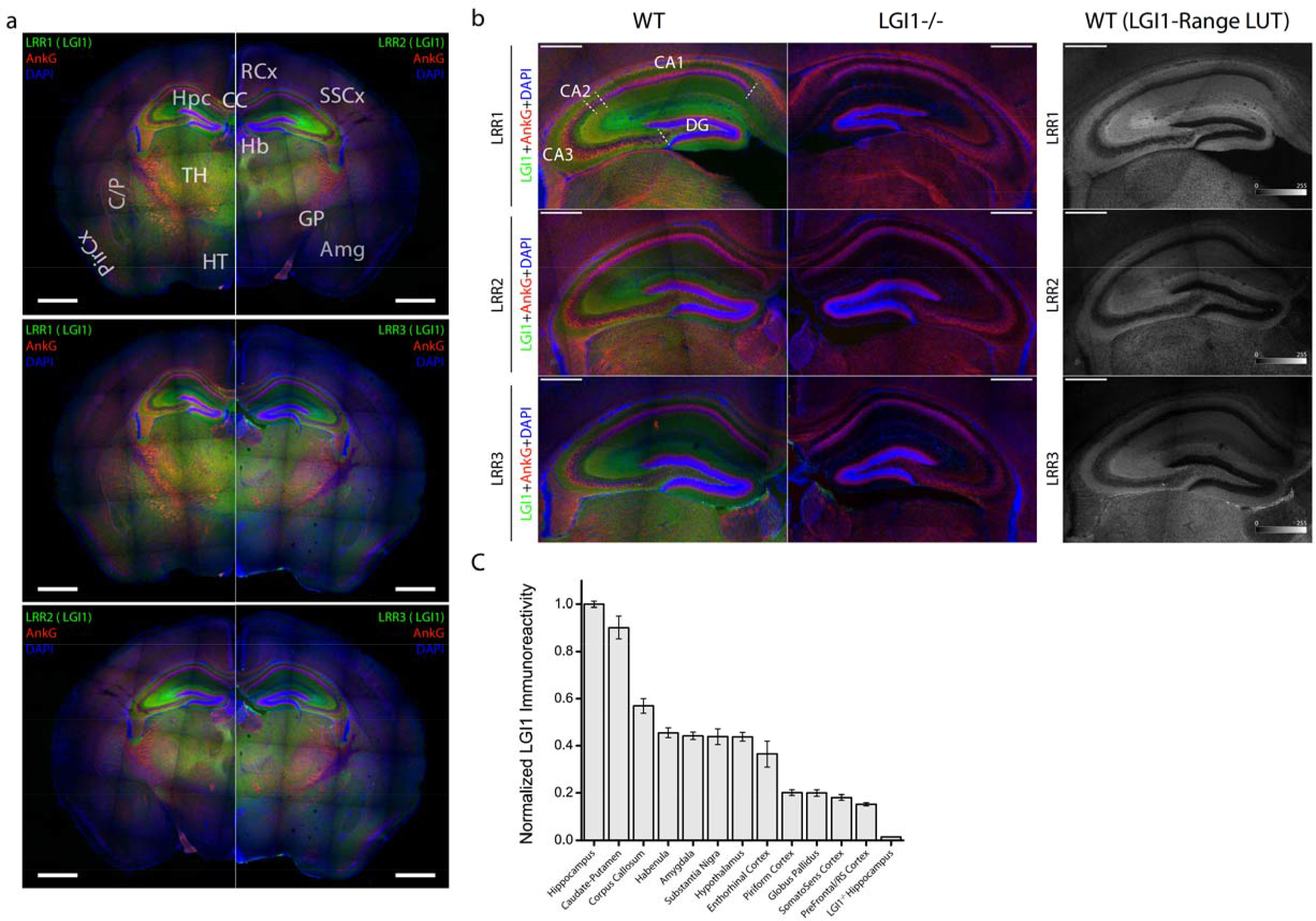
Validation and characterization of human monoclonal anti-LGI1 antibodies in the mouse brain. **a** Paired comparison of the immunoreactivity pattern of different antiobies in the mouse hippocampus showing LRR1 vs. LRR2 (top panels), LRR1 vs. LRR3 (middle panels), and LRR2 vs. LRR3 (bottom panels). The LGI1 signal is always shown in green. Slices were counter-stained with AnkG (red) and DAPI (blue). Scale bar = 1 mm. **b** Antibody validation showing a complete absence of immunostaining in the LGI1^-/-^ hippocampus for LRR1 (top panels), LRR2 (middle panels) and LRR3 (bottom panels). In the right column, a range-color look-up table has been assigned to the LGI1 signal of each one of the antibodies to visually compare their immunostaining patterns. Scale bar = 0.5 mm. **c** Comparison of normalized immunoreactivity levels of LGI1 in different prosencephalic and mesencephalic structures. Columns and error bars show mean ± SEM. n = 4 independent experiments (WT vs *Lgi1^-/-^*; LRR1 n = 1; LRR2 n = 2; LRR3 n = 1). n Hippocampus=90 ROIs, n Amygdala = 90 ROIs; n Corpus callosum = 65 ROIs; n Caudate/Putamen = 55 ROIs; n Globus Pallidus = 30 ROIs; n Substantia nigra = 60 ROIs; n Habenula = 60 ROIs; n Hypothalamus = 95 ROIs; n Entorhinal cortex = 25 ROIs, n Piriform Cortex= 110 ROIs; n Prefrontal cortex = 160 ROIs; n Somatosensory Cortex = 95 ROIs; n *Lgi1^-/-^* hippocampus = 135 ROIs. Data obtained with different antibodies were normalized to WT hippocampal levels in each experiment, and pooled thereafter

**Fig. 2.**
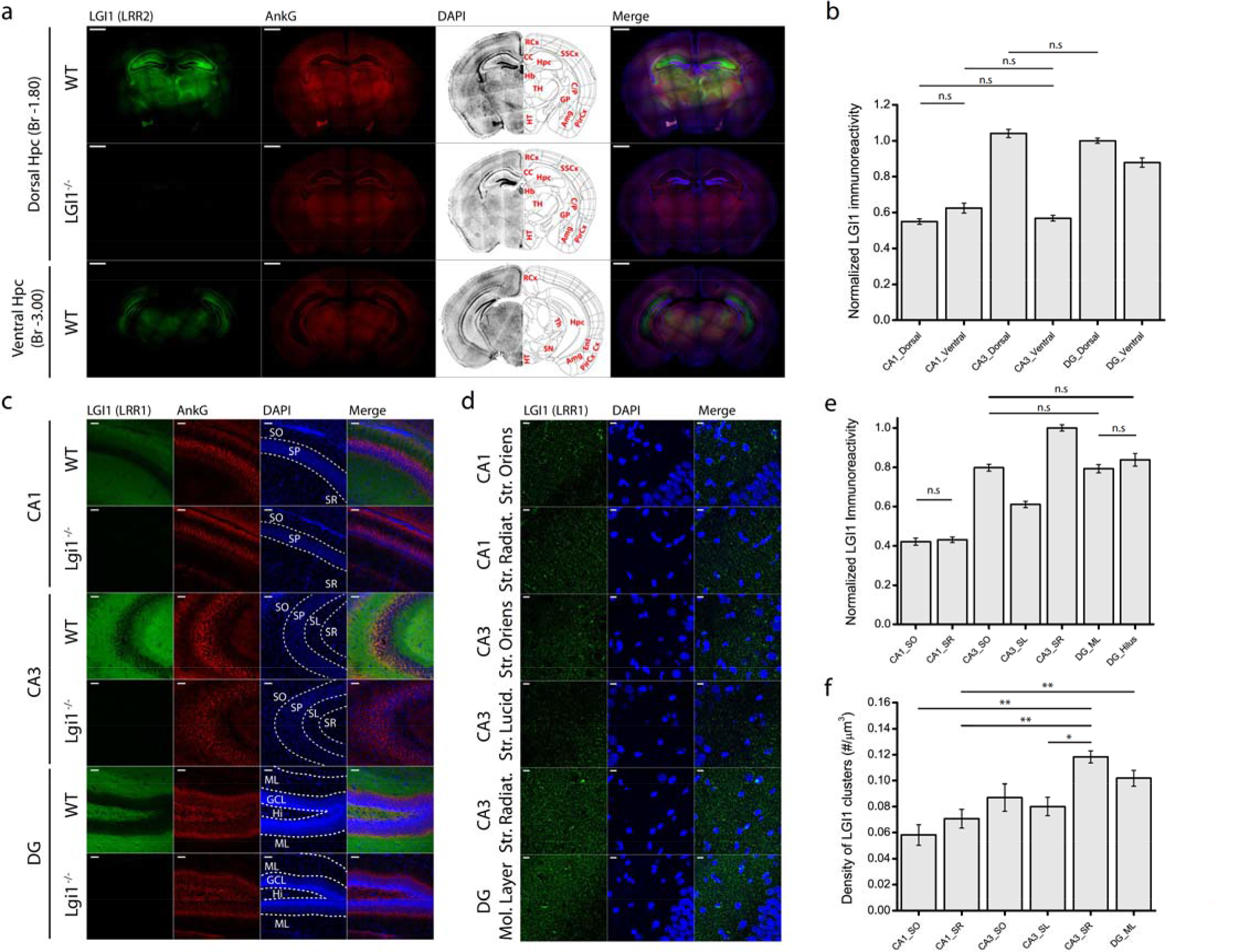
Characterization of the hippocampal pattern of expression and distribution of LGI1. **a** LGI1 (LRR2) immunostaining in different rostro-caudal levels through the mouse brain showing a differential expression of LGI1 in the dorsal and the ventral hippocampus. WT dorsal hippocampus (Bregma approx. -1.80mm, top panels). LGI1^-/-^ dorsal hippocampus (Bregma approx. -1.80mm, middle panels). WT ventral hippocampus (Bregma approx. -3.00mm, top panels). LGI1 signal is shown in green, AnkG in red, and DAPI in blue. Scale bar = 1mm. **b** Quantification of LGI1 expression at different hippocampal levels. Columns and error bars show mean ± SEM. n=4 animals (LRR1 n = 1; LRR2 n = 2; LRR3 n = 1). For the sake of clarity only non-significant comparisons are shown. Any other comparison is \*\**p*<0.01. n CA1 Dorsal=140 ROIs; n CA1 Ventral = 110 ROIs; n CA3 Dorsal = 210 ROIs; n CA3 ventral = 195 ROIs; n DG Dorsal = 210 ROIs; n DG Ventral = 165 ROIs. Data obtained with different antibodies were normalized to DG Dorsal levels in each experiment and pooled thereafter. **c** LGI1 (LRR1) immunostaining throughout different dorsal hippocampal regions showing specific immunolabelling of LGI1 in CA1 (top panels), CA3 (middle panels), and DG (bottom panels). LGI1 signal is shown in green, AnkG is in red, and DAPI is in blue. Scale bar = 50 µm. **d** High-magnification pictures showing a punctate pattern of staining with anti-LGI1 (LRR1) in the different hippocampal strata analysed. LGI1 is shown in green, and DAPI is in blue. Scale bar = 10 µm**. e** Quantification of LGI1 expression in different hippocampal subregions within the dorsal hippocampus. Columns and error bars show mean ± SEM. N = 4 animals (LRR1 n = 1; LRR2 n = 2; LRR3 n = 1). For the sake of clarity only non-significant comparisons are shown. Any other comparison is \*\**p*<0.01. n CA1 Str. Oriens = 70 ROIs; n CA1 Str. Radiatum = 70 ROIs; n CA3 Str. Oriens = 70 ROIs; n CA3 Str. Lucidum = 70 ROIs; n CA3 Str. Radiatum = 70 ROIs; n DG Molecular Layer = 140 ROIs; n DG Hilum = 70 ROIs. Data obtained with different antibodies was normalized to CA3 Stratum Radiatum levels in each experiment and pooled thereafter. **f** Quantification of LGI1 puncta densities (number of clusters/µm^3^) in the different hippocampal strata within the dorsal hippocampus. Columns and error bars show mean ± SEM. n=2 independent experiments (WT vs *Lgi1^-/-^*, LRR1 n=1; LRR2 n=1). For the sake of clarity only significant comparisons are shown. \**p*<0.05; \*\**p*<0.01. Any other comparison is non-significant. n CA1 Str. Oriens = 8 Fields of view; ROIs; n CA1 Str. Radiatum = 8 Fields of view; n CA3 Str. Oriens = 8 Fields of view; n CA3 Str. Lucidum = 8 Fields of view; n CA3 Str. Radiatum = 8 Fields of view; n DG Molecular Layer = 8 Fields of view

A closer observation of the stratum oriens (SO) of both CA1 and CA3, as well as the DG granule cell layer, showed that LGI1 clusters are often attached to ankyrin-G-dense axonal initial segments (AIS) (Fig. 3). This is well-demonstrated in the three-dimensional reconstruction of AIS (Fig. 3b) with a CA3 >DG > CA1 gradient of LGI1 expression (Fig. 3c). In order to address whether LGI1 staining in CA3_SL and CA3_SR corresponded to postsynaptic dendritic staining of CA3 pyramidal neurons or to presynaptically expressed LGI1, we co-stained both LGI1 and the presynaptic synaptic vesicle marker synaptophysin. Partial colocalization between LGI1 and synaptophysin was observed in all hippocampal strata (Online Resource 1) with a slight but significant reduction of this colocalization seen in CA3_SL when compared to CA3_SR or DG_ML (Online Resource 1b). The strong immunoreactivity of CA3_SR and the partial co-localization with synaptophysin suggest LGI1 staining in the CA3_SR is, at least partially, attributable to the dendritic expression of LGI1 in CA3 pyramidal cells. Therefore, we evaluated the colocalization of LGI1 with the postsynaptic marker PSD93 in CA1, CA3 and DG layers. As shown in Online Ressources 1a and 2a, Manders’ overlap coefficients (Online Resource 1b and 2b) indicate a limited however more prominent LGI1::PSD93 colocalization index compared to LGI1::synaptophysin. This finding corroborates the presence of a significant postsynaptic LGI1 pool.

**Fig. 3.**
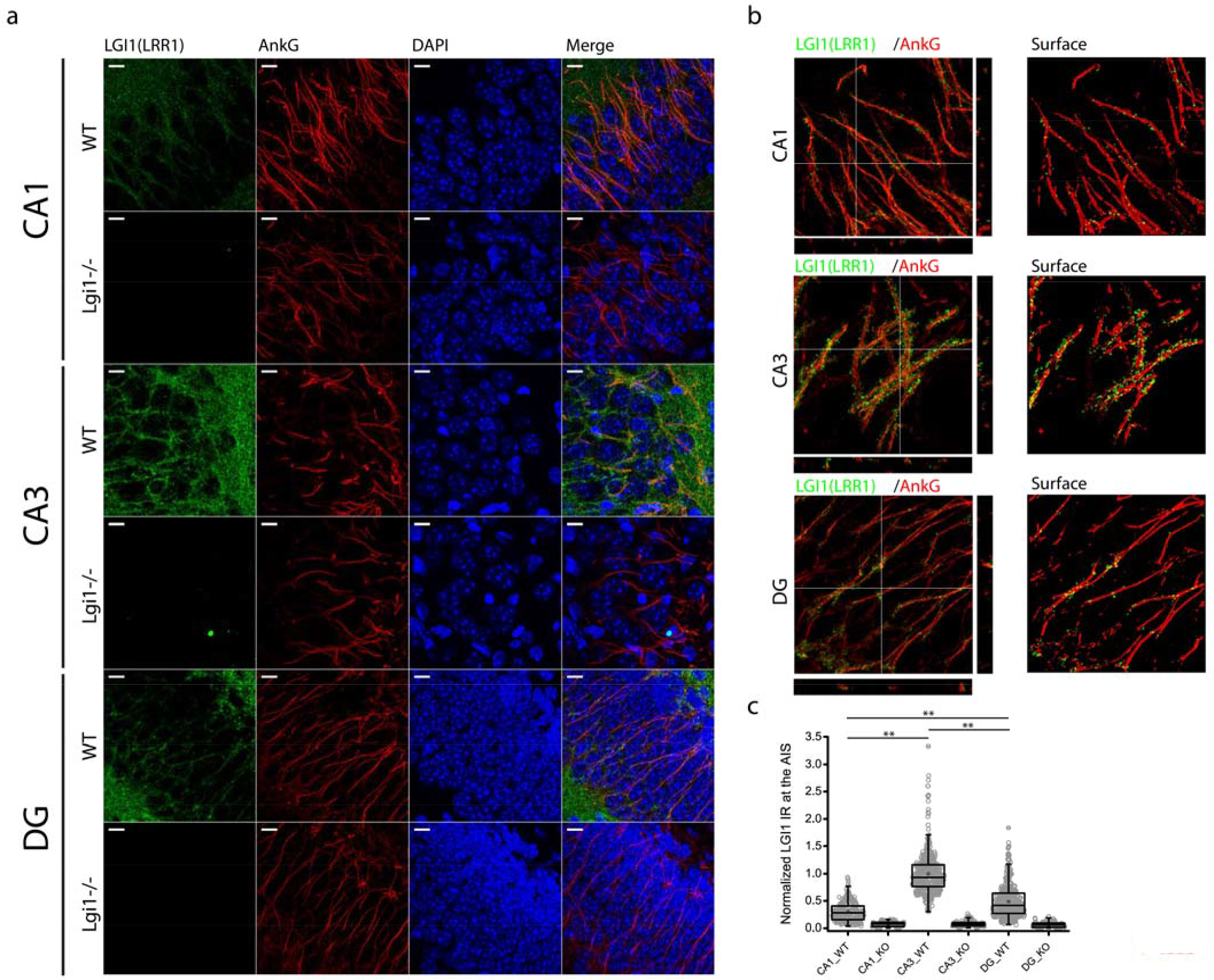
LGI1 is differentially expressed at the AIS of distinct hippocampal neurons. **a** High-magnification pictures showing specific LGI1 (LRR1) immunostaining at the AIS of CA1 pyramidal cells (top panels), CA3 pyramidal cells (middle panels), and DG granule cells (bottom panels). LGI1 signal is shown in green, AnkG is in red, and DAPI is in blue. Scale bar = 10 µm. **b** Orthogonal views (left columns) and surface reconstruction (right columns) of LGI1 signal (LRR1, green) restricted to the AIS (AnkG, red) of CA1 pyramidal cells (top panels), CA3 pyramidal cells (middle panels), and DG granule cells (bottom panels). **c** Quantification of LGI1 immunoreactivity at the AIS of different hippocampal neuronal subtypes. n = 2 independent experiments (WT vs *Lgi1^-/-^*; LRR1 n = 1; LRR2 n = 1). For the sake of clarity only comparisons within the WT genotype are shown. \**p*<0.05; \*\**p*<0.01. n CA1 WT= 260 AIS; n CA1 *Lgi1^-/-^* =126 AIS; n CA3 WT = 425 AIS; n CA3 *Lgi1^-/-^* = 156 AIS; n DG WT = 427 AIS; n DG *Lgi1^-/-^* = 191 AIS. Data obtained with different antibodies were normalized to CA3-AIS levels in each experiment, and pooled thereafter

### Astrocytic expression of LGI1

While LGI1 has been almost exclusively considered as a trans-synaptic protein, it was first described in astrocytic gliomas. Also, strong LGI1 mRNA expression was reported in astrocytes (The human protein atlas database; www.proteinatlas.org). Therefore, we investigated the expression of LGI1 in hippocampal astrocytes with a focus on the CA3 hippocampal subregion (Fig. 4a and b). As shown in Fig. 4a (CA3) and b (CA3_SO), GFAP-expressing astrocytes in CA3 mainly populate CA3_SO and CA3_SR and, consequently, are present in the CA3 strata enriched for LGI1. Given the widespread punctate staining of LGI1 in all CA3 strata, maximal projections of the z-stacks required to fully cover the span of astrocytes (>20 optical slices; 0.38 µm/slice) were inappropriate to resolve the precise localization of LGI1 within astrocytic processes (Fig. 4c, top). Reduced z-stacks (3 optical slices which focus on the span of specific astrocytic processes) show LGI1 immunoreactive puncta that can be ascribed to some of the single astrocytic processes (Fig 4c, bottom). Also, AnkG co-staining showed occasional astrocytic juxtaposition to the AIS of CA3 pyramidal neurons, forming astrocyte-AIS contact points (Figs. 4b and d). While its sporadic nature ruled out a detailed quantitative analysis, in the CA3_SO, LGI1 was consistently observed at astrocytes-AIS contact points (Fig. 4e and f).

**Fig. 4.**
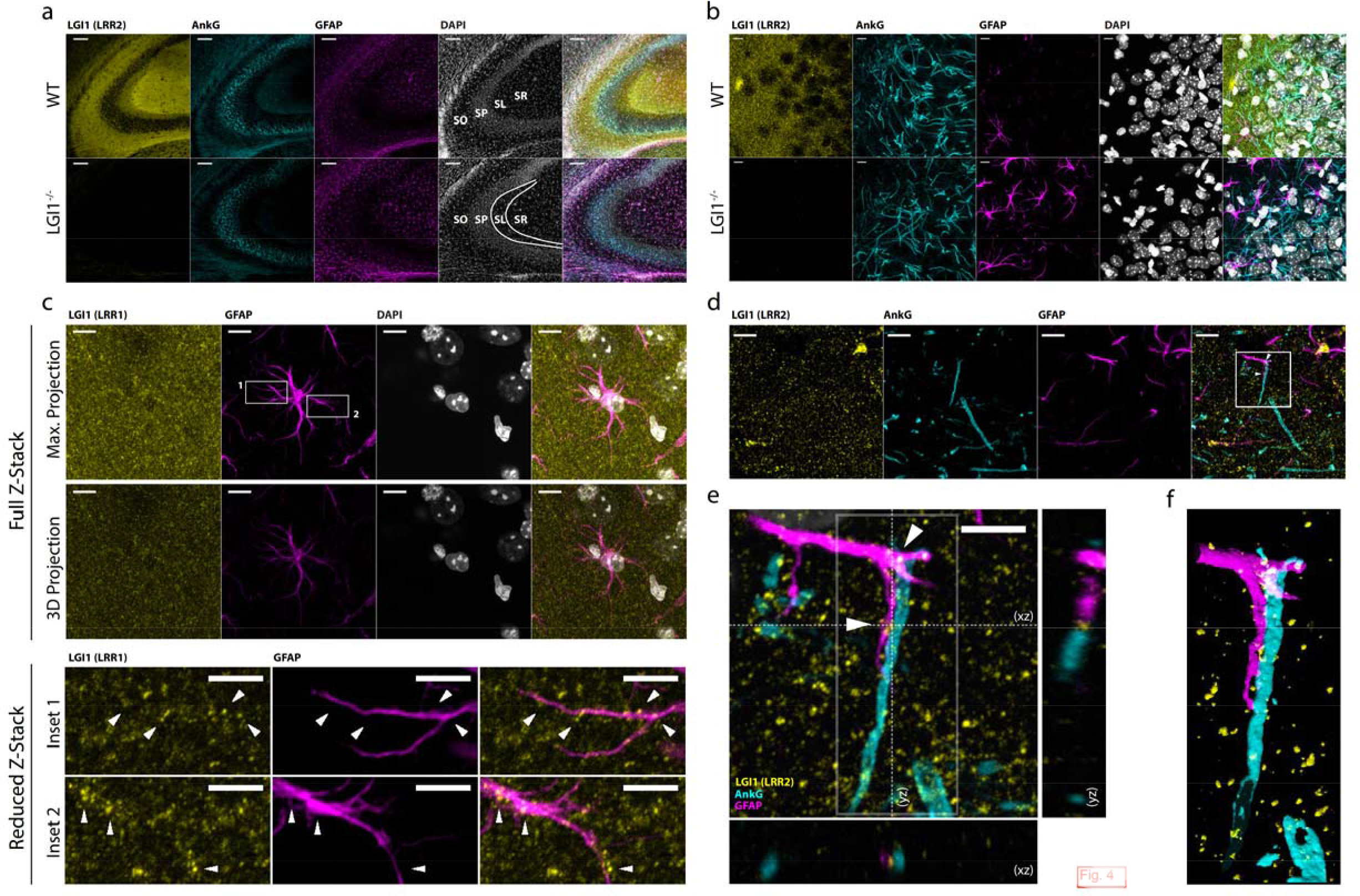
LGI1 is expressed by hippocampal astrocytes. **a, b** LGI1 (LRR2, yellow), AnkG (cyan), GFAP (magenta), and DAPI (gray) staining in CA3 mouse hippocampus at low magnification (**a**) and at high magnification (**b**) in the Stratum Oriens of CA3. Scale bars = 100 µm (**a**) and 10 µm (**b**). **c** Maximal projections and 3D projections of a Z-stack spanning a CA3-hippocampal astrocyte showing LGI1 (LRR1, yellow), GFAP (magenta), and DAPI (gray) staining (top) and reduced Z-stack of the two astrocytic processes corresponding to the insets (1, 2) outlined in the top rows (bottom). Arrowheads indicate LGI1 puncta over the astrocytic processes. Scale bars = 5 µm. **d** LGI1 (LRR2, yellow), AnkG (cyan), GFAP (magenta), and DAPI (gray) staining in the Stratum Oriens of CA3 showing an astrocytic process in close contact with an axon initial segment stained with GFAP and AnkG, respectively. **e** Orthogonal views of the region boxed in **d** showing that LGI1 (LRR2, yellow) is precisely located between the astrocytic process (GFAP, magenta) and the axonal initial segment (AnkG, cyan). Scale bar = 5 µm. **f** Volume reconstruction of the region boxed in **e**. Images are representative of 3 independent experiments (WT vs *Lgi1^-/-^*; LRR2 n = 2; LRR1 n = 1)

### Subcellular distribution of secreted LGI1

To observe LGI1 secretion from neurons, we transfected by Gene Gun rat hippocampal organotypic slices with Dendra2-fused LGI1 (D2::LGI1), and used an anti-dendra2 antibody to visualise intracellular and secreted LGI1 5-7 days post-transfection (Fig. 5a). LGI1 was observed across neuronal arborisations (Fig. 5b), including the axon, AIS and dendrites of the CA1 (Fig. 5c), CA3 (Fig. 5d) and DG (Fig. 5e) transfected neurons. In the DG, a differential subregional expression was observed within dendrites ≥ AIS > axon (Fig. 5e). The AIS LGI1 punctae were often observed in AnkG gap-like microdomains (Fig. 5f), previously shown to express GABAaα2 receptor clusters at axo-axonal synapses [38] [79]. Indeed, we also identified GABAaα2 subunits adjacent to LGI1 (Fig. 5g).

**Fig. 5.**
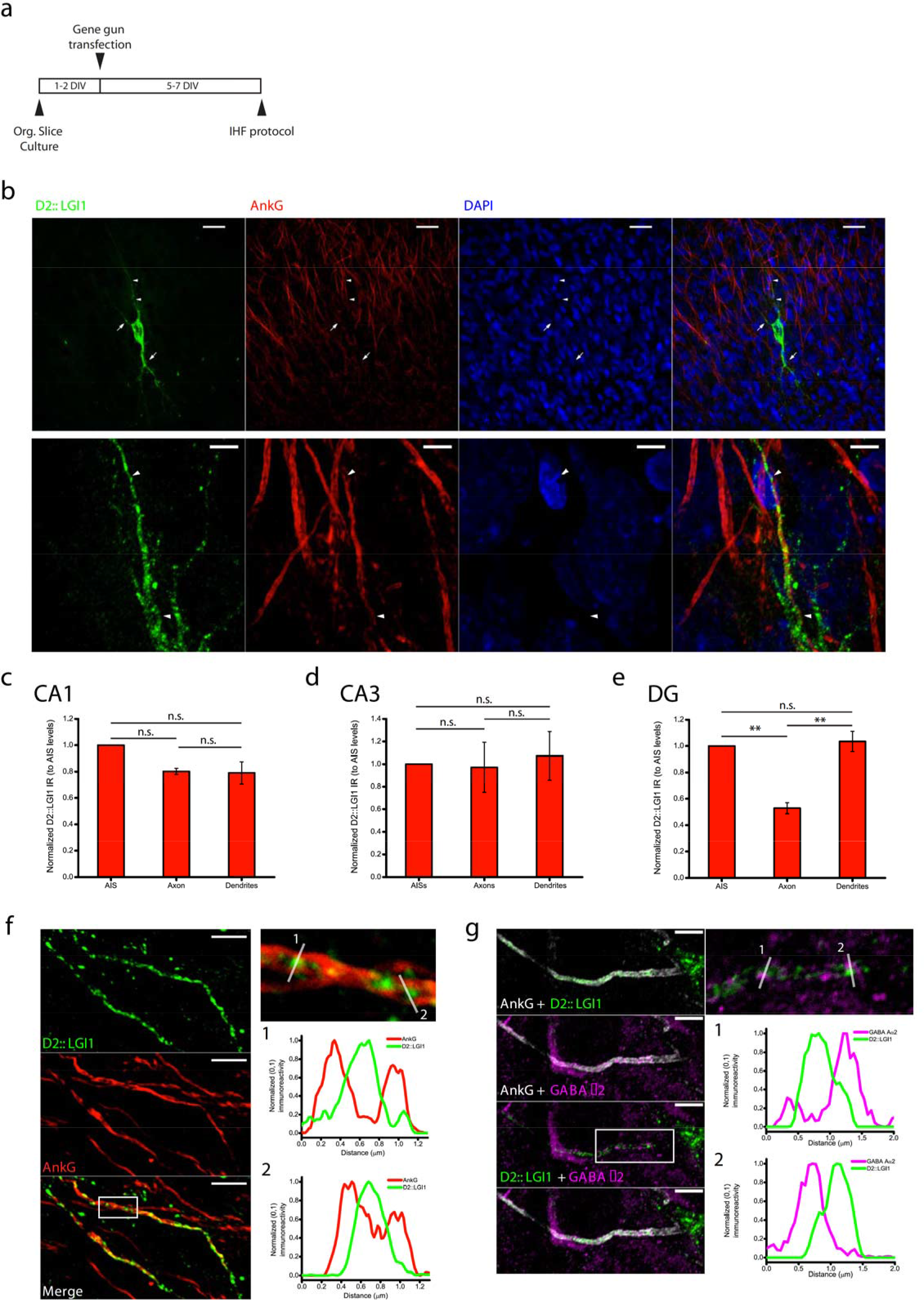
Subcellular distribution of total DENDRA2::LGI1 studied by gene-gun transfection and immunohistofluorescence in rat organotypic slices. **a** Schematic of the experimental protocol used for gene-gun transfection and immunostaining. **b** Top panels, CA3 pyramidal cells transfected with DENDRA2::LGI1 and labelled with anti-DENDRA2 (green), anti-AnkG (red), and DAPI (blue). Arrowheads point towards the axonal initial segment. Arrows show dendritic branches. Scale bar = 25 µm. Bottom panels, high-magnification picture showing the expression of LGI1 at axonal initial segment (arrowheads) and surrounding dendrites. Scale bar = 5 µm. **c, d, e** Quantification of the relative immunoreactivity levels of DENDRA2::LGI1 at the AIS, axon, and dendrites of transfected neurons in CA1 (**c**), CA3 (**d**), and DG (**e**). Columns and error bars show mean ± SEM. n = 2 independent organotypic cultures. n CA1 = 3 cells; n CA3 = 3 cells; n DG = 5 cells. n.s. non-significant; * *p*<0.05; \*\**p*<0.01. **f** Left columns, High-magnification picture of two axonal initial segments from two transfected cells showing prominent expression of DENDRA2::LGI1 at the AIS. DENDRA2::LGI1 is shown in green, AnkG is in red. Scale bar = 5 µm. Right columns, detail of the outlined inset showing that DENDRA2::LGI1 is partially distributed in the AnkG gap-like microdomains at the AIS (top panel). Normalized immunoreactivity levels for DENDRA2::LGI1 and AnkG (green and red traces, respectively) of the lines depicted in the inset (bottom panels). **g** Immunostaining of DENDRA2::LGI1 (green), AnkG (gray), and GABA Aα2 (magenta), at the AIS of a CA3 transfected neuron (top panels). Scale bar = 5 µm. Traces for DENDRA2::LGI1 and GABA Aα2 normalized immunoreactivity (green and magenta, respectively) of the lines shown in the inset (bottom panels)

Next, as Gene Gun transfection can be stochastic, we electroporated D2-tagged (D2::LGI1) LGI1 into selected CA3 pyramidal neurons in organotypic slices to dissect LGI1 distribution versus secretion in CA3 neurons. The identical appearances of D2::LGI1 distribution in mice and rat cultures (Online Ressource 3) justified use of this system to profile secreted D2::LGI1 by applying anti-dendra antibodies to live cultures before fixation. As shown in Fig. 6b, this procedure robustly labelled punctae of secreted LGI1 surrounding the electroporated neuron, around the cell body, proximal apical dendrites, the AIS and a distal axonal region of the transfected neurons (Fig. 6c). Several D2::LGI clusters colocalized with PSD93 at the AIS of transfected neurons (Fig. 7 and Online Resource 4), as well as in thorny excrescences and proximal apical dendrites of CA3 pyramidal neurons (Online Resource 4). Colocalisation of secreted D2::LGI1 with PSD93 was similar in all analysed regions of the transfected neurons (Fig. 7c,d).

**Fig. 6.**
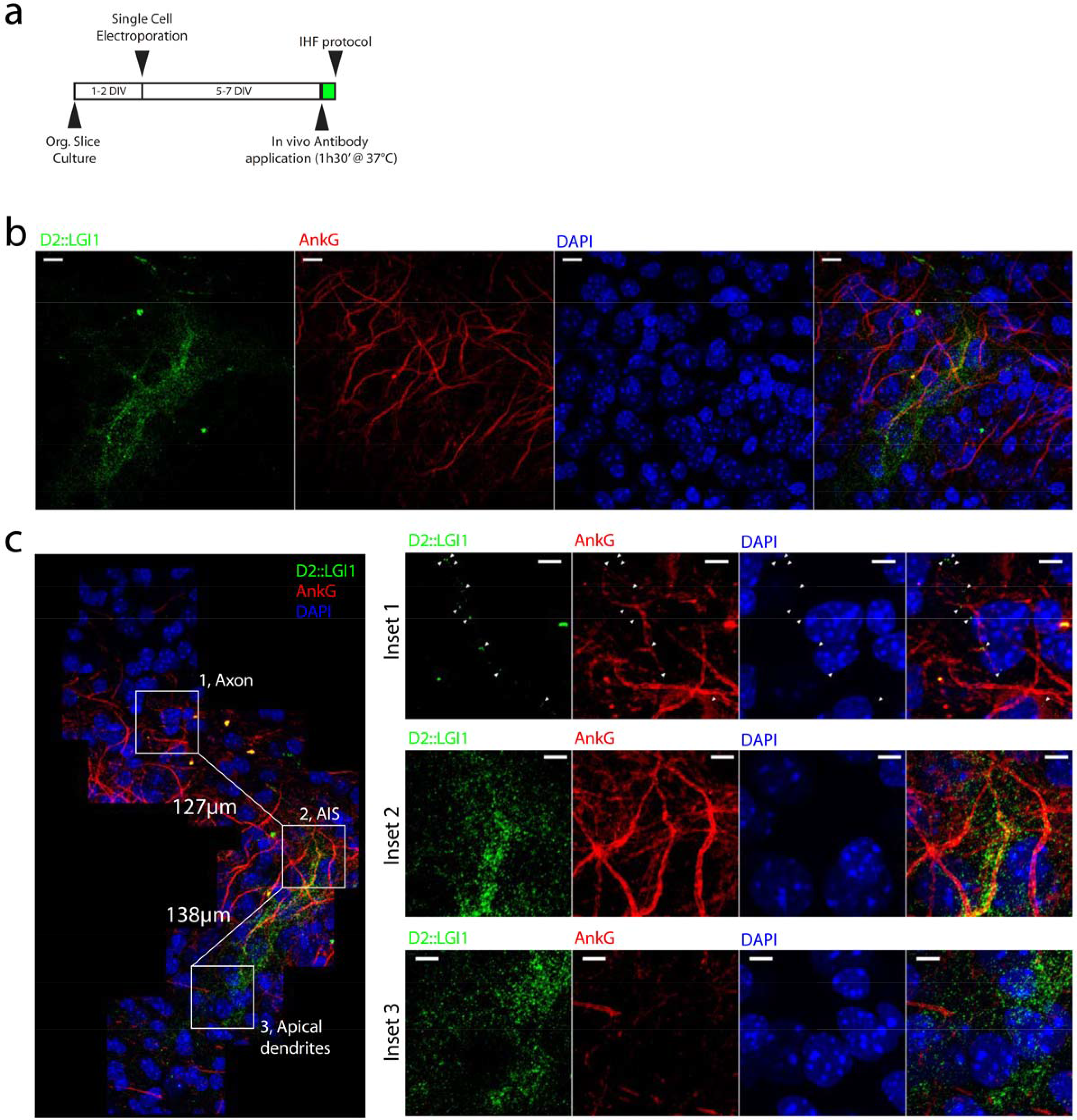
DENDRA2::LGI1 is reliably secreted by CA3 pyramidal cells after single cell electroporation. **a** Schematic of the experimental protocol used for single cell electroporation and immunostaining for extracellular detection of DENDRA2::LGI1. **b** Extracellular (secreted) DENDRA2::LGI1 revealed by anti- DENDRA2 (green) in an electroporated CA3 pyramidal cell. AnkG is shown in red, and DAPI is in blue. Scale bar = 10 µm. **c** High-magnification reconstruction of the electroporated neuron shown in (**b**). Outlined insets are shown in the right panels where inset 1 corresponds to the axon (arrowheads indicate axonal secretion spots of DENDRA2::LGI1), inset 2 correspond to the axonal initial segment, and inset 3 corresponds to the proximal apical dendrite. Images are representative of 3 CA3 pyramidal neurons from 2 independent cultures. Scale bars = 5 µm in all panels

**Fig. 7.**
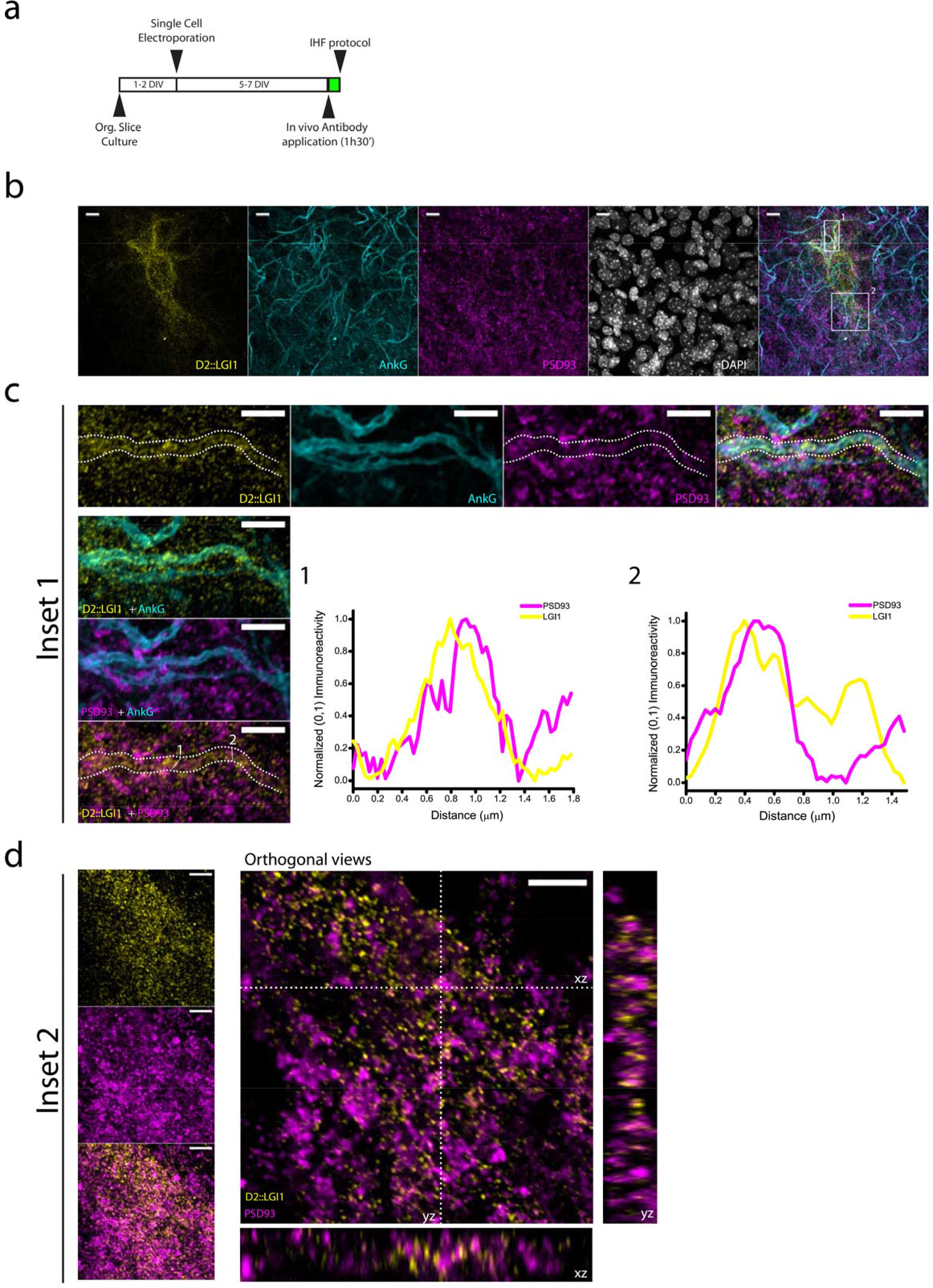
Secreted DENDRA2::LGI1 colocalizes with AnkG and PSD93 after single cell electroporation in mouse hippocampal slices. **a** Schematic of the experimental protocol used for single cell electroporation and immunostaining. **b** Low-magnification picture of two individual CA3 pyramidal neurons electroporated with DENDRA2::LGI1 and immunostained against secreted DENDRA2::LGI1 with anti- DENDRA2 (yellow), anti-AnkG (cyan), anti-PSD93 (magenta), and DAPI (gray). Scale bar = 10 µm. **c** High magnification detail of the inset 1 outlined in panel **b** showing colocalization of secreted DENDRA2::LGI1 (yellow) with the AIS marker AnkG (cyan), and PSD93 clusters (magenta) at the AIS. Dashed lines indicate the edges of the AIS taking AnkG signal as a reference. Line traces (1,2) correspond to the line traces depicted in the bottom left panel. Scale bar = 5 µm. **d** High magnification detail of the inset 2 outlined in panel **b** showing colocalization of secreted DENDRA2::LGI1 (yellow) with PSD93 clusters (magenta) at the dendrites of CA3 pyramidal neurons. Maximal projections (left panels) and orthogonal views (right panels) of the basal dendrites of CA3 pyramidal neurons shown in **b**. Scale bars = 5 µm

To confirm the specificity of this distribution, we electroporated and followed untagged LGI1 immunostaining in *Lgi1^-/-^* slices, using LRR1. The observed subcellular distribution of LGI1 closely resembled that of D2::LGI1 (Fig. 8, inset 1) with LGI1 clusters colocalized with AnkG in the AIS, and also in dendritic branches and at the tip of individual dendritic spines (Fig. 8, Inset 2). These findings confirm an important cis-enrichment of LGI1 in dendrites and postsynaptic compartments.

**Fig. 8.**
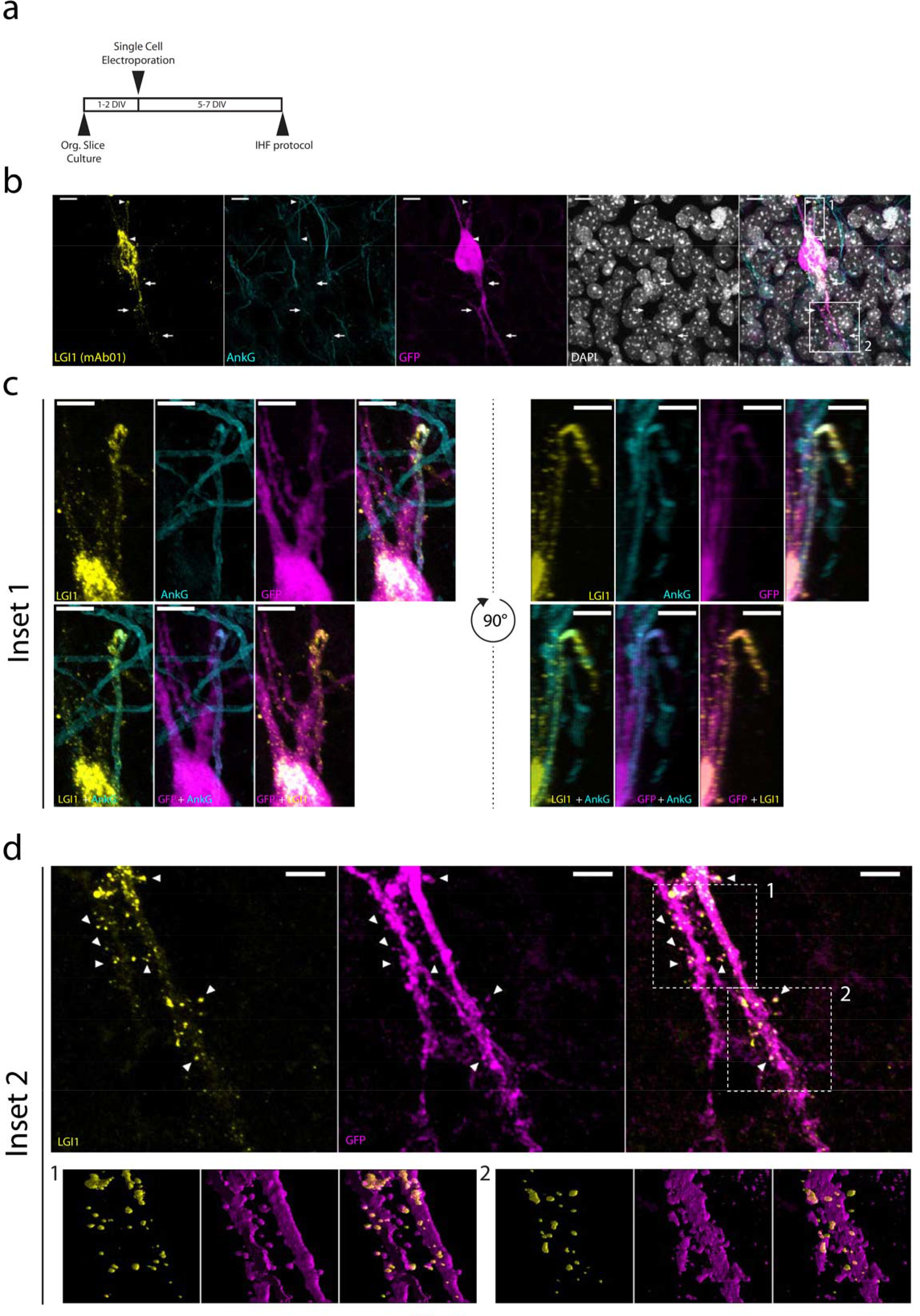
WT LGI1 localizes to the AIS and dendrites of CA3 pyramidal neurons after single cell electroporation in mouse *Lgi1^-/-^* hippocampal slices. **a** Schematic of the experimental protocol used for single cell electroporation and immunostaining. **b** CA3 pyramidal cell transfected with WT LGI1 and soluble GFP and labelled with anti-LGI1 (Yellow), anti-AnkG (cyan), anti-GFP (magenta) and DAPI (grey). Scale bars = 10 µm. **c** Detailed view of the region outlined in b (1) corresponding to the AIS of the electroporated neuron showing the appearance of LGI1 clusters at the AIS (left panels) and 90° rotated view (right panels). Scale bars = 5 µm. **d** Detailed view of the region outlined in b (2) showing LGI1 clusters at the proximal basal dendrites of the electroporated neurons. Arrowheads point towards individual dendritic spines in which LGI1 clusters were identified (top panels). Volume reconstruction of the regions 1 and 2 boxed in the top panels (bottom panels). Scale bars = 5 µm

### LGI1-associated proteome

In order to identify protein complexes in which native LGI1 is expressed and, at the molecular level, address the potential heterogeneity in LGI1-associated protein complexes suggested by its immunolocalization, we performed immunoprecipitation and mass spectrometry analysis of wild type and *Lgi1^-/-^* rodent brain homogenates using the non-competing antibodies (LRR1 & 3 [63]) and also EPTP1 & 2. EPTP1 & 2 specifically recognized LGI1-EPTP domain (Online Resource 5 a, b, c) and do not recognize overlapping epitopes (Online Resource 5 d). Full string networks for the LGI1 interactome associated with all antibodies were generated using string-db.org and are represented in Online Ressources 7, 8, 9, 10 & 11. As expected, LGI1 was efficiently recovered by all antibodies (Fig. 9 and Spreadsheet). Each antibody showed a specific interactome (Fig. 9, Table 1 and Spreadsheet) with common partners immunoprecipitated by either two (Table 2) or three different antibodies (Table 3). Our data confirm several previously reported proteins such as ADAM proteins, pre and postsynaptic scaffolds as well as potassium channel subunits [39] [21] [67] and also uncovers several novel partnerships, showing the existence of different LGI1 complexes selectively identified by different patient-derived antibodies. These partners are principally neuronal but glial-associated proteins were also identified and their subcellular distributions are presented hereafter within a cell-specific context.

**Fig. 9.**
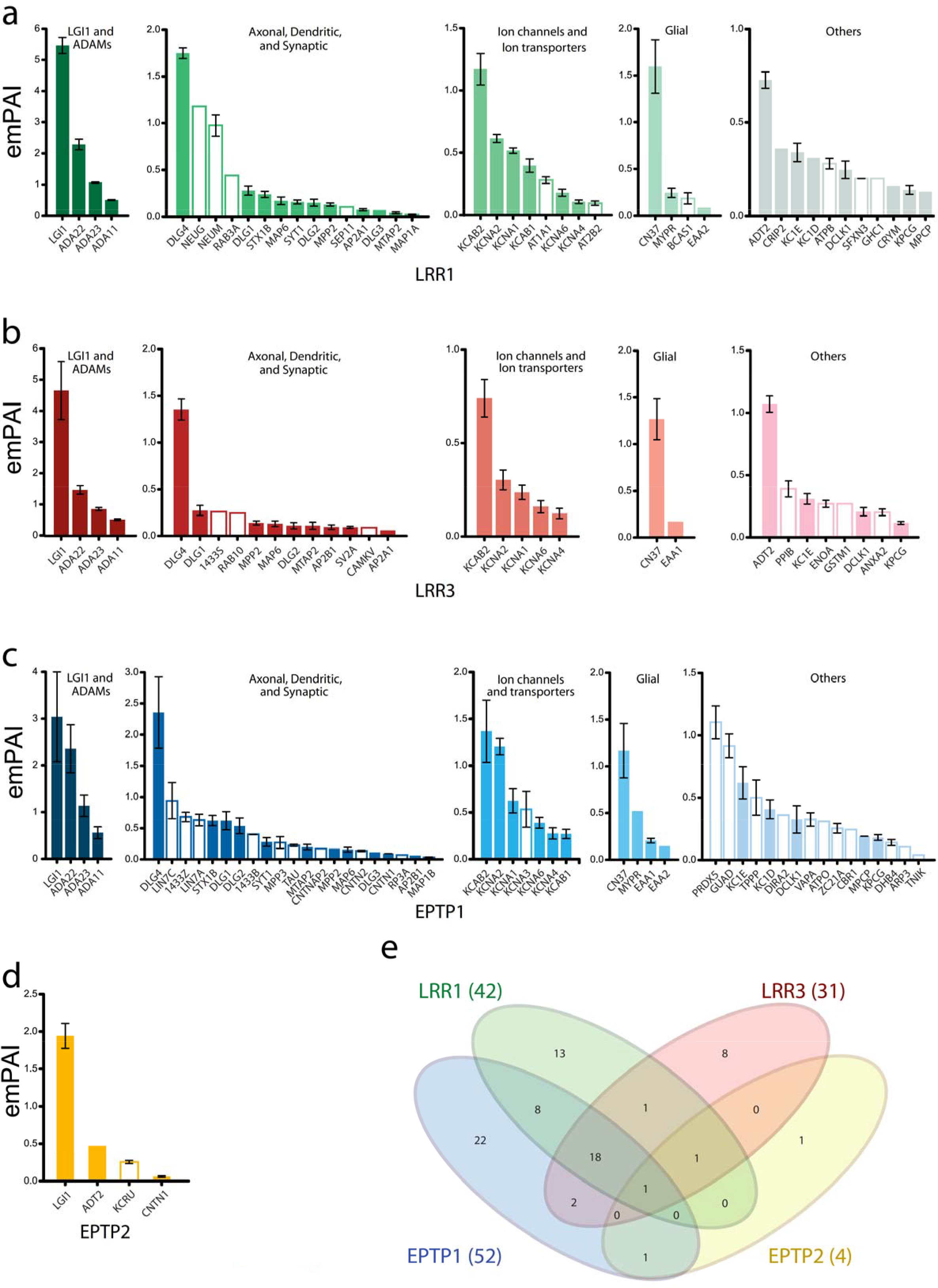
Histograms of specific LGI1 partners immunoprecipitated by the recombinant patient-derived anti- LGI1 antibodies. Absolute mean emPAI values of LGI1 partners immunoprecipitated by LRR1 (a), LRR3 (b), EPTP1 (c) and EPTP2 (d) illustrates their relative enrichment. Partners exclusively immunoprecipitated by a specific antibody are represented by open histograms. For clarity, tubulins are not represented in these histograms and are listed in Spreadsheet. A Venn Diagram (d) shows the number of overlapping LGI1 partners immunoprecipitated by the antibodies used. Immunoprecipitated partners are classified by functional classes (Partially from http://www.pantherdb.org/)

**Table 1.** LGI1 partners specific to each IgG emPAI of immunoprecipitated partners are listed. *associated with lipid rafts; # Present in myelin sheaths proteome

| Antibody | Gene Symbol | Description |
| --- | --- | --- |
| **LRR1** | AT1A1 (*)<br>AT2B2<br>ATPB (*)<br>BCAS1 (#)<br>CRIP2<br>CRYM<br>GHC1<br>MAP1A (*)<br>NEUG<br>NEUM (*) (#)<br>RAB3A (#)<br>SEP11 (#)<br>SFXN3 (#) | Sodium/potassium-transporting ATPase subunit alpha-1<br>Plasma membrane calcium-transporting ATPase 2<br>ATP synthase subunit beta, mitochondrial<br>Breast carcinoma-amplified sequence 1 homolog<br>Cysteine-rich protein 2<br>Ketimine reductase mu-crystallin<br>Mitochondrial glutamate carrier 1<br>Microtubule-associated protein 1<br>Neurogranin<br>Neuromodulin<br>Ras-related protein Rab-3A<br>Septin-11<br>Sideroflexin-3 |
| **LRR3** | 1433S (#)<br>ANXA2 (#)<br>CAMKV<br>ENOA<br>GSTM1 (#)<br>PPIB<br>RAB10<br>SV2A | 14-3-3 protein sigma<br>Annexin A2<br>CaM kinase-like vesicle-associated protein<br>Alpha-enolase<br>Glutathione S-transferase Mu 1<br>Peptidyl-prolyl cis-trans isomerase B<br>Ras-related protein Rab-10<br>Synaptic vesicle glycoprotein 2A |
| **EPTP1** | 1433B (*) (#)<br>1433Z (#)<br>ARP3<br>ATPO<br>CBR1<br>CNTN2 (*)<br>CNTNAP2<br>DHB4<br>DIRA2<br>GUAD<br>KCNA3 (#)<br>LIN7A<br>LIN7C<br>MAP1B (#)<br>MPP3<br>PRDX5<br>RP3A<br>TAU<br>TNIK<br>TPPP (#)<br>VAPA<br>ZC21A | 14-3-3 protein beta/alpha<br>14-3-3 protein zeta/delta<br>Actin-related protein 3<br>ATP synthase subunit O, mitochondrial<br>Carbonyl reductase [NADPH] 1<br>Contactin-2<br>Contactin-associated protein-like 2<br>Peroxisomal multifunctional enzyme type 2<br>GTP-binding protein Di-Ras2<br>Guanine deaminase<br>Potassium voltage-gated channel subfamily A member 3<br>Protein lin-7 homolog A<br>Protein lin-7 homolog C<br>Microtubule-associated protein 1B<br>MAGUK p55 subfamily member 3<br>Peroxiredoxin-5, mitochondrial<br>Rabphilin-3A<br>Microtubule-associated protein tau<br>Traf2 and NCK-interacting protein kinase<br>Tubulin polymerization-promoting protein<br>Vesicle-associated membrane protein-associated protein A<br>Zinc finger C2HC domain-containing protein 1A |
| **EPTP2** | KCRU | Creatine kinase U-type, mitochondrial |

(\*) Present in lipid rafts proteome (#) Present in myelin sheaths proteome

**Table 2.** LGI1 partners immunoprecipitated by two distinct antibodies emPAI of immunoprecipitated partners are listed. *associated with lipid rafts; # Present in myelin sheaths proteome

| Antibody | Gene Symbol | Description |
| --- | --- | --- |
| LRR1<br>∩<br>EPTP1 | DLG3 (SAP102)<br>EAA2<br>KC1D<br>KCAB1<br>MPCP<br>MYPR (*) (#)<br>STX1B (*)<br>SYT1 (*) (#) | Disks large homolog 3<br>Excitatory amino acid transporter 2<br>Casein kinase I isoform delta<br>Voltage-gated potassium channel subunit beta-1<br>Phosphate carrier protein, mitochondrial<br>Myelin proteolipid protein<br>Syntaxin-1B<br>Synaptotagmin-1 |
| LRR3<br>∩<br>EPTP1 | AP2B1<br>EAA1 | AP-2 complex subunit beta<br>Excitatory amino acid transporter 1 |
| LRR1 ∩ LRR3 | AP2A1 | AP-2 complex subunit alpha-1 |
| EPTP2 ∩ EPTP1 | CNTN1 ((*) (#) | Contactin-1 |

(\*) Present in Lipid rafts proteome (#) Present in myelin sheaths proteome

**Table 3.** LGI1 partners immunoprecipitated by three antibodies emPAIs of immunoprecipaitated partners are listed. *associated with lipid rafts; # Present in myelin sheaths proteome.

| Gene Symbol | Description |
| --- | --- |
| ADA11 | Disintegrin and metalloproteinase domain-containing protein 11 |
| ADA22 | Disintegrin and metalloproteinase domain-containing protein 22 |
| ADA23 (#) | Disintegrin and metalloproteinase domain-containing protein 23 |
| CN37 (*) (#) | 2',3'-cyclic-nucleotide 3'-phosphodiesterase |
| DCLK1 | Serine/threonine-protein kinase DCLK1 |
| DLG1 (SAP97) | Disks large homolog 1 |
| DLG2 (PSD93) | Disks large homolog 2 |
| DLG4 (PSD95) (*) | Disks large homolog 4 |
| KC1E | Casein kinase I isoform epsilon |
| KCAB2 (#) | Voltage-gated potassium channel subunit beta-2 |
| KCNA1 (#) | Potassium voltage-gated channel subfamily A member 1 |
| KCNA2 (#) | Potassium voltage-gated channel subfamily A member 2 |
| KCNA4 | Potassium voltage-gated channel subfamily A member 4 |
| KCNA6 | Potassium voltage-gated channel subfamily A member 6 |
| KPCG | Protein kinase C gamma type |
| MAP6 (#) | Microtubule-associated protein 6 |
| MPP2 | MAGUK p55 subfamily member 2 |
| MTAP2 | Microtubule-associated protein 2 |
| ADT2 (*) | ADP/ATP translocase 2 |

(\*) Present in Lipid rafts proteome

### Neurons: axonal, dendritic and synaptic

Our immunofluorescence results indicate that LGI1 is not targeted to a specific neuronal compartment, however its significant colocalization with PSD93 suggests a relative enrichment at synaptic contact sites. Besides previously recognized partners such as ADAM11, 22 and 23, Kv1 channel subunits and pre/post-synaptic MAGUKs, LRR3 immunoprecipitated the calcium-dependent phospholipid-binding protein annexin 2 (ANXA2) that is known to be associated, in GABAergic interneurons [84], with the extracellular leaflet of plasma membranes at cell-cell contact sites [85]. LRR1 also recognised LGI1- associated complexes in axon-terminals, since it efficiently immunoprecipitated neuromodulin (GAP43). This palmitoylated phosphoprotein is involved in synaptic function modulation [57] and is enriched in axon terminals, where it regulates actin cytoskeleton dynamics and participates in signalling pathways by promoting PI(4.5)P2 clustering in cholesterol rich domains [9]. Presynaptic complexes were highlighted by the efficient immunoprecipitation by EPTP1 of the effector protein rabphilin-3A, the presynaptic CASK-binding scaffolding proteins Lin7A (Veli1) and 7C (Veli3). A unique link with G-protein signalling is also highlighted by this antibody since it co-immunoprecipitated LANC2 (LanC-like protein 2), a myristoylated peripheral phosphoinositide-phosphate binding protein that activates adenylate cyclase through interaction with G-proteins [17]. The strong staining of LGI1 in dendritic structures is corroborated by a significant association with neurogranin and with several postsynaptic scaffold proteins such as PSD93, PSD95, SAP97 and SAP102 (Table 3, Online Resource 6 and Spreadsheet). The postsynaptic MTAP2 as well as the axonal STOP protein MAP6 [29] were also efficiently retained by LRR1, LRR3 and EPTP1 confirming the immunofluorescent identification of LGI1 in stable dendritic and axonal compartments.

### Cytoskeletal

Immunoprecipitation of microtubule-associated proteins (LRR1, LRR3 and EPTP1) (Fig. 9 and Table 3), tubulins (LRR1 and EPTP1) (Table 2) and the axonal marker tau (EPTP1) (Table 1 and Spreadsheet) as well as the cytoskeletal GTPase Septin-11 show that, although intra-vesicular, LGI1 coimmunoprecipitates with cytoskeletal proteins most likely via vesicle membrane-partners [40].

### Axo-glial junctions

EPTP antibodies selectively captured potential juxtaparanodal protein complexes. EPTP1 immunoprecipitated contactin 1 (CNTN1), TAG1 (CNTN2) and Caspr2 (CNTNAP2) while EPTP2 coimmunoprecipitated only CNTN1.

### Neurons: vesicular trafficking

LRR1, LRR3 and EPTP1 highlighted common and distinct links of LGI1 to vesicular trafficking pathways. While LRR1 co-immunoprecipitated Rab3a, an effector of exocytosis implicated in fine tuning synaptic vesicle release and synaptic plasticity [42] [15], LRR3 was associated with synaptic vesicle glycoprotein 2A (SV2A) known to be particularly involved in GABAergic synaptic transmission [55]. EPTP1 was the only IgG that enriched VAPA, (Vesicle-associated membrane protein-associated protein A. This protein mediates ER / plasma membrane contact sites [36] through its interaction with oxysterol-binding protein-related protein 3 and is involved in several cellular functions such as lipid transport, Kv2 exocytosis, membrane trafficking and microtubule reorganisation [73]. AP2 complex subunits as well as syntaxin1 and synaptotagmin1 were co-immunoprecipitated by several antibodies (Table 2).

### Kinases and other enzymes

Alongside the neuron-specific PKCγ (KPCG) [66] immunoprecipitated by LRR1, LRR3 and EPTP1 (Table 3), several kinases such as casein kinase 1ε, 1δ, doublecortin-like kinase 1 (DCLK1) and other enzymes were also immunoprecipitated by different LGI1 antibodies (Tables 1, 2 & 3). This is in line with the fact that several phosphorylation pathways are involved in Kv1 targeting [78] and that Kv1 expression is perturbed in *Lgi1^-/-^* mice.

### Rafts

Despite the diversity of the observed protein complexes, all antibodies recovered proteins previously described as part of the lipid raft proteome [35] (Tables 1, 2 and 3). LRR1, 3 and EPTP2 recovered a considerable amount of the antiporter ADP/ATP translocase 2 (ADT2 / SLC25A5), a master regulator of mitochondrial energy enriched in microglia and endothelial cells [2]. LRR1 immunoprecipitated neuromodulin (GAP-43), a major component of growth cones involved in long term potentiation and synaptic function [57]. Other proteins enriched in lipid rafts were immunoprecipitated by several antibodies like tubulin beta subunits, microtubule associated protein 1A as well as the Na/K ATPase subunit α1 and the ATP synthase β subunit of the mitochondrial ATP synthase (Spreadsheet). Other lipid raft-enriched partners with specific characteristics are mentioned within the corresponding compartmental classification.

### Oligodendrocytes

All LGI1-antibodies immunoprecipitated proteins known to be part of the myelin proteome [32] (Spreadsheet and Tables 1, 2 & 3). LRR1, LRR3 and EPTP1 uncovered a strong and specific enrichment of CN37 (2’,3’-cyclic-nucleotide 3’-phosphodiesterase), a marker for non-myelinating oligodendrocytes [3] [44] (Table 3). LRR1 selectively captured Bcas1, which defines an oligodendrocytic population involved in active myelin formation [13]. EPTP1 also enriched tubulin polymerization-promoting protein (TPPP), a crucial actor in myelin sheath elongation in oligodendrocytes [18].

### Astrocytes

A limited number of partners suggesting an astrocytic association were recovered. EPTP1 immunoprecipitated the astrocytic excitatory amino acid transporters EAA1 (GLAST-1) and EAA2 (GLT-1) and LRR3 specifically immunoprecipitated Glutathione S-transferase Mu 1 (GSTM1) implicated in proinflammatory astrocyte-microglia communication [37].

### Other

Partly because of their ubiquitous expression, the enrichment of some proteins was not straightforward to interpret. Alpha-enolase (ENOA1), stratifin (14-3-3 S) and peptidyl-prolyl cis-trans isomerase B (PPIB) were recovered by LRR3 and the mitochondrial proteins creatine kinase KCRU and amino acid transporter GHC1 (Slc25a22) were recovered by EPTP2 and LRR1 respectively (Fig. 9 and Table 1). It is interesting however to mention that 14-3-3 proteins were reported to stabilize LGI1/ADAM22 interaction [82] and that ENO1 was recently shown to protect against cerebral ischemia-induced neuronal injury [33]. Although specific, the co-association of the plasma membrane calcium-transporting ATPase2 (AT2B2) is very weak and awaits further confirmation.

## Discussion

In this study, we used a set of recombinant LGI1-autoantibodies, derived from two patients with LGI1- antibody limbic encephalitis, to address the precise localisation of native LGI1 and its interactome. Our immunofluorescent and immunoprecipitation data converge, expanding the known synaptic localisation of LGI1 to include somatodendritic and axonal subcellular localisations and to highlight diverse oligodendrocytic, neuro-oligodendrocytic and astro-microglial complexes containing LGI1. These, and the cytoskeletal, vesicular, enzyme and membrane raft-enriched proteins complexed with LGI1, provide focussed pathways to study LGI1’s functions in basic neurobiology and in the context of multiple human diseases.

Three LRR-antibodies (LRR1, LRR2 and LRR3) were used on fixed brain or organotypic slices to investigate the detailed distribution of native LGI1 throughout mice forebrain and hippocampal structures, and to follow its secretion profile. Both LRR- and EPTP-antibodies were used to dissect the composition of LGI1-associated protein complexes. Two of the three antibodies (LRR1 and 2) used for immunostaining have overlapping epitopes [63] which do not overlap with LRR3. All three antibodies gave very similar signals and were specific with respect to non-staining of *Lgi1^-/-^* tissue. We found that LGI1 is differentially expressed in brain regions with sub-regional profiles that correlate with observed dysregulations in LGI1-antibody limbic encephalitis patients. These include the hippocampal focal CA3 atrophy in these patients; abnormal caudate and putamen imaging often observed in the ∼60% of LGI1- antibody patients who have faciobrachial dystonic seizures; and the serum hyponatraemia which may relate to dysregulation of hypothalamic structures [31] [16].

LGI1 is predominantly expressed in hippocampus, thalamus and hypothalamus with a lower expression in the cortical and subcortical regions. In the hippocampus, the CA3 and the DG were the most intensely labelled regions and LGI1 was equally distributed in somatodendritic and axonal projections. Interestingly, normalized immunoreactivities indicated that the CA3_SR was the most intensely labelled zone. This region is highly interconnected, receiving perforant-path inputs. Mossy fibers synapse onto local inhibitory interneurons in CA3_SR and onto the proximal apical dendrites of excitatory pyramidal cells at thorny excrescences in CA3_SL. In CA1, CA3 and DG, LGI1 immunoreactive puncta were highly enriched at axonal initial segments (AIS), and partially overlapped with the synaptic vesicle marker synaptophysin. The presence of LGI1 at the AIS is in agreement with the observed increase in intrinsic excitability in *Lgi1 ^-/-^* mice [83] [68] [86]. Also, the presynaptic effects in *Lgi1^-/-^* mice [83] [5] or in an LGI1 secretion mutant [87] are in agreement with the observed presynaptic expression and immunoprecipitated complexes. Although pre- and postsynaptic effects in *Lgi1^-/-^* mice have been reported [83], exclusive pre- or postsynaptic effects have also been described. Reported postsynaptic effects in *Lgi1 ^-/-^* mice [21] [54] [46] are in agreement with the dendritic expression of this protein and the presence of postsynaptic LGI1-associated protein complexes. Considering the prominent expression of LGI1 in the DG, the reduced staining of LGI1 in CA3_SL and its limited co-localization with synaptophysin, our data imply LGI1 is not solely enriched in mossy fibres from the granule cells but may have a major postsynaptic localisation. We corroborated this by showing a significant colocalization with the postsynaptic marker PSD93.

LGI1 association with vesicular trafficking pathways and the enrichment of VAPA by EPTP1 as well as the presence of LGI1 at Ankyrin G deficient regions adjacent to GABAα2 clusters at the AIS suggest that LGI1 may be important in the organization and dynamics of neuronal signalling pathways [38] that contribute to control of AIS excitability. LRR1, LRR3 and EPTP1 immunoprecipitation profiles highlighted common and distinct links of LGI1 to vesicular trafficking pathways. While LRR1 co-immunoprecipitated Rab3a, an effector of exocytosis implicated in fine tuning synaptic vesicle release and synaptic plasticity, LRR3 was associated with synaptic vesicle glycoprotein 2A (SV2A) known to be particularly involved in GABAergic synaptic transmission. EPTP1 was the only IgG that enriched VAPA, (Vesicle-associated membrane protein-associated protein A. This protein mediates ER / plasma membrane contact sites through its interaction with oxysterol-binding protein-related protein 3 and is involved in several cellular functions such as lipid transport, Kv2 exocytosis, membrane trafficking and microtubule reorganisation. AP2 complex subunits as well as syntaxin1 and synaptotagmin1 were co-immunoprecipitated by several LGI1 antibodies (Table 2).

Fluorescent recombinant LGI1 transfection in CA3 pyramidal neurons allowed us to address its secretion profile. LGI1 did not show preferential exocytic sites and was found to be similarly secreted from axonal and somatodendritic regions. Of note, in addition to its expression at the AIS, somatodendritic expression of LGI1 was also reported in mature cultured hippocampal neurons [27]. The diffuse distribution of LGI1 in native tissue suggests that depending on its localisation, LGI1 participates in multiple distinct functional protein complexes and that its role may not be restricted to trans-synaptic complexes. While no major structural changes in neurons were identified in *Lgi1 ^-/-^* mice [5], an increase in dendritic arborization and a decrease in dendritic pruning were reported [87]. However, no aberrant dendritic sprouting or synaptic pruning defects were observed in mice infused with anti-LGI1 [58]. This may indicate that despite traces of neuronal damage in LGI1-dependent LE patients [43], either some LGI1 protein complexes are not accessible to patient-derived autoantibodies or that despite the presence of antibodies, LGI1 molecules are still able to modulate dendritic pruning. Also, LGI1 involvement in dendritic sprouting and synaptic pruning may be purely developmental and thus not observed during antibody application in mature animals.

In contrast to previous reports of relatively restricted LGI1-associated complexes [21] [39], mass spectrometry analysis of immunoprecipitated proteins uncovered an intriguingly rich and complex LGI1 interactome, which converged on specific cell types and pathways. Together with GFAP co-labelling, the presence of LGI1 (of neuronal or astrocytic origin) at neuron-astrocyte contact points and the enrichment of juxtaparanodal protein complexes (contactin1, TAG1 and Caspr2) with some LGI1-antibodies, may have functional implications for its role in glial biology [14] [1]. Although we cannot conclude that LGI1 is systematically enriched in hippocampal astrocytes, some astrocytic processes clearly show the presence of immunoreactive LGI1 clusters. An interaction of LGI1 with astrocytic cell adhesion molecules [6] cannot be excluded and needs further investigation. The neuronal and oligodendrocytic GPI-anchored TAG1 is a crucial determinant of axo-glial interactions [23] and is involved in Caspr2 and Kv1 channel complexes at juxtaparanodes [77]. It is also enriched with Kv1.2 in the axonal initial segment [59]. The single transmembrane domain protein Caspr2 is an axonal protein that interacts extracellularly with TAG1 and binds MPP2 and CASK [28] that were co-immunoprecipitated in this study. These observations may concur with phenotypic overlaps observed in patients who have Caspr2-directed antibodies [62] and provide a molecular link to hypomyelination reported in *Lgi1^-/-^* mice [71]. Our data also suggest that Caspr2/ADAM22/23 complexes [27] as well as LGI1/ADAM22/23 occur in association with Kv1 at the AIS and/or glutamatergic synapses. The constant presence of ADAM proteins, MAGUKS and Kv1 subunits in the interactomes is consistent with the hypothesis that LGI1-antibodies destabilize membrane Kv1 channels leading to hyperexcitability, however the precise mechanisms involved await elucidation. Interestingly, the glial protein CNPase was unambiguously identified by three distinct antibodies however, the molecular link mediating LGI1 interaction with this multifunctional intracellular protein [60] deserves additional investigation.

Although our data do not allow us to determine to what extent LGI1 participates in trans synaptic complexes, the observed localisation of native LGI1 as well as the diversity of LGI1-linked complexes are not in favour of a major implication of LGI1 as trans synaptic linker proteins. Of particular note, no AMPA or NMDA receptor-associated subunits were immunoprecipitated with any of the autoantibodies. This suggests that, if such interactions resist our mild solubilisation conditions, AMPAR/NMDAR/LGI1 complexes may not be recognized by the LGI1-antibodies or that all our antibodies specifically dissociated from these complexes. These assumptions seem unlikely since three of the antibodies efficiently enriched proteins containing PDZ domains, especially PSD95 (DLG4), known to independently co-associate with ADAM22 [19] and AMPA receptor subunits [11]. Indeed, it is interesting to note that LGI1 was not reported in an AMPA receptor interactome, nor AMPARs in LGI1 complexes [70] [21] [39]. A decrease in total and synaptic Kv1.1 precedes AMPA receptor downregulation in mice infused with patient-derived whole LGI1-IgGs [58]. Taken together, these results question a molecular link between LGI1 and AMPA receptors. To reconcile these findings with the literature, the observed downregulation of these postsynaptic receptors in LGI1-linked pathologies may be a homeostatic response to hyperexcitability induced by a reduction in Kv1 expression [68] [58]. Additional research is needed to clarify this issue.

Consistent with the hypothesis that anti-LGI1 antibodies may initiate Kv1 channel downregulation, we identified Kv1.1, 1.2, 1.4 and 1.6 subunits as well as Kv beta subunits (KCAB1 & 2) associated with LGI1, in agreement with previous reports [21]. Presumably ADAM22/11 and MAGUKS provide the predominant link between Kv1 and LGI1. A direct association between ADAM23 and Kv1 is unlikely since ADAM23 lacks a PDZ interaction domain [49]. LGI1 mRNA expression levels in astrocytes, neurons, microglial cells and oligodendrocytes show an important heterogeneity (www.proteinatlas.org) and therefore, providing a relatively stable mRNA/protein ratio and similar secretion profile from one cell type to the other, the abundance of certain LGI1 associated protein complexes may differentially depend on the nature of each antibody and the LGI1 secretion point.

In conclusion, our immunostaining and mass spectrometry results indicate strong regional expression of LGI1 that is consistent with imaging of hippocampal damage in patients with LGI1 limbic encephalitis. Subcellular localisation supports a role for LGI1 in regulating intrinsic excitability and synaptic transmission. Our data reconcile a large number of contradictory reports regarding the neurobiology of LGI1, notably by demonstrating LGI1 secretion in both axonal and somatodendritic compartments and highlighting LGI1 expression at glial contact sites and its association with glial proteins. Our findings support the hypothesis that LGI1 antibodies in patients with limbic encephalitis induce hyperexcitability by destabilizing Kv1 channels, notably at axonal initial segments. However, the precise mechanisms by which these antibodies trigger downregulation require further investigation.

## Acknowledgements

This work was supported by the Institut National pour la Science et la Recherche Médicale (INSERM), Aix-Marseille Université (AMU) and the Agence Nationale de la Recherche (ANR) (grant ANR-17-CE16- 0022). The postdoctoral financial support of J.R.F. and the PhD thesis support of J.E. were from the ANR (grant ANR-17-CE16-0022). The PhD thesis of K.D. was supported by a fellowship from the French Ministry of Research (MESRI). The facility for recombinant protein production in eukaryotic cell systems of the AFMB laboratory is supported by the French Infrastructure for Integrated Structural Biology (FRISBI) (grant ANR-10-INSB-05-01). Acquisition of the mass spectrometer was financially supported by the “Fédération de Recherche pour le Cerveau” (FRC) through the Rotary operation “Espoir en tête”. We thank Stephanie Baulac for sharing the Lgi1^-/-^ mice [7] and the pcDNA3.1-FLAG-LGI1 plasmid. We also thank Stephen Elledge for providing the pINDUCER11 (miR-RUG) plasmid. We are grateful for UCB Pharma, 208-216 Bath Road, Berkshire, UK who generated and purified human recombinant monoclonal anti-LGI1 IgGs. We thank Faivre-Sarrailh for critically reading the manuscript.

## Declarations

### Contributions

OEF conceived the study, supervised the entire project, the experimental design, data interpretation and manuscript preparation. OEF and JRF designed the study, analysed and interpreted the data. JRF performed immunofluorescence experiments on slices, images treatments and analysis. SP performed immunofluorescence experiments on HEK cells and corresponding images treatments. JRF performed Excel-based mass spectrometry analysis. LEF performed Python-based scripts for mass spectrometry analysis. JE performed brain slicing, organotypic cultures and single cell electroporation experiments. KD performed immunoprecipitation experiments and participated in immunofluorescence experiments and data analysis. JRF, KD and JE took care of mice breeding and availability of Lgi1^-/-^ animals. MB and CV performed mass spectrometry experiments. CD and PM generated His-tagged LGI1 expressing viral particles, transfected cultured Sf9/21 cells and provided filtered conditioned culture medium. YM purified His-tagged recombinant LGI1. CL performed SPR experiments. DD and MR supervised the work of JE. SI provided human monoclonal anti LGI1 antibodies from limbic encephalitis patients. YM, YF and MS performed expression plasmids preparation and preliminary expression tests of recombinant constructs in heterologous systems. MS and CL contributed to the design of the study. OEF and JRF wrote the original draft of the manuscript. OEF and JRF prepared the figures. All authors edited and reviewed the manuscript.

### Competing interests

The authors declare that they have no competing interests.

### Data and material availability

Data supporting this manuscript are available in the Electronic Supplementary Materials.

## Electronic supplementary material

### Supplementary Material and Methods

#### Construction of plasmids for inducible LGI1 expression in mammalian-expression

pIND-IRES-EGFP comprises the Tet inducible promoter region and the entire expression cassette under the control of the ubiquitin promoter. The 473 bp Tet inducible promoter of pINDUCER11 was amplified with the forward (<u>CAATTCAGTCGACT**GGATCC**TTTA</u>CC) and reverse (GATCGA**GCGGCCGC**CAGTGTGATGGATATC<u>T</u>GCAGAATTCAGGCTGGAT<u>CG</u>GTCCCG) primers (underlined are overlap sequences with pINDUCER11), digested with BamHI and NotI (in bold) and inserted into pcDNA3.1 pre-digested with BglII/NotI. The resulting plasmid was digested with DraIII and ligated with MluI adaptors, then digested with BstBI and ligated with PstI adaptors. This 3539 bp vector fragment was ligated with the 4003 bp MluI/PstI fragment of pINDUCER11, yielding pIND-IRES-EGFP vector (7556 bp).

pIND-LGI1-IRES-EGFP was constructed as follows: the LGI1 sequence was obtained from pCDNA3.1-FLAG-LGI1 [7]. LGI1 sequence was PCR-amplified and modified by nested PCR, introducing an optimized Kozak consensus sequence in 5’ and removing the FLAG tag. The signal peptide region was amplified with the forward (GGTACCGAGCTCGGACGAATTCCACC<u>ATGGAATCAGAAAGCAGCAGAAG</u>G) and reverse (GGCATTTTGGCTTCGCTGGTTTC<u>T</u>TCCCCT<u>CAGTC</u>AGC) primers (underlined are overlaps with the LGI1 signal peptide coding sequence). The secretory LGI1 sequence was amplified with the forward (GCTGACTGAGGGGAAGA<u>A</u>AC<u>CAGCGAAGCCAAAATG</u>CC) and reverse BGHR (TAGAAGGCACAGTCGAGG) primers (underlined is overlapping region with the LGI1 coding sequence). Both amplification products were mixed in equimolar amounts and amplified with the external (GGTACCGAGCTCGGACGAATTCCACC<u>ATGGAATCAGAAAGCAGCAGAAG</u>G) and BGHR (TAGAAGGCACAGTCGAGG) primers. The resulting 1775 bp LGI1 insert was digested with EcoRI / NotI and ligated into the same sites of pIND-IRES-EGFP vector.

pIND-ΔIRES-Dendra2-LGI1: pIND-ΔIRES (6204 bp) lacking the IRES-EGFP module, was first obtained by AscI / PacI digestion of pIND-IRES-EGFP, blunting and re-ligation. Dendra2 coding sequence was inserted downstream of LGI1 signal-peptide coding sequence. It was obtained by PCR from pDendra2-LifeAct7 (Addgene #54694) and cloned by the SLIC method [2] using the forward <u>CTGACTGAG</u>GGGAAGAAGATGAACACCCCGGGAATTAAC and reverse primers <u>TTTGGCTTCGCTG</u>GTGAACCTCCAGACCACACCTGGCTGG (underlined are overlaps with LGI1 sequence).

#### Antibodies

Rabbit (386003, RRID: AB_2661876) and guinea pig (386005, RRID: AB_2737033) anti- ankyrinG (AnkG) and rabbit anti-GFP (132003, RRID: AB_1834147) were from Synaptic Systems. Mouse anti-synaptophysin 171B5 was a generous gift from Dr. S. Fujita, Mitsubishi Kasei Institute, Japan. Mouse anti-GFAP (75-240, RRID: AB_10672299) and mouse anti PSD- 93 (75-057, RRID: AB_2277296) were from Antibodies Incorporated. Rabbit anti-DENDRA2 (ABIN361314, RRID: AB_10789591) was from Antibodies-Online. Alexa coupled 488-goat anti-human and anti-rabbit, 594-goat anti-mouse, anti-rabbit and anti-mouse, 647-goat anti-mouse and anti-guinea pig were from Jackson ImmunoResearch.

#### Immunohistofluorescence of fixed brains

P14-P16 C57BL/6 wild-type mice or *Lgi1*^-/-^ littermates of either sex were deeply euthanized with a Ketamine-Xylazine mix (100mg/Kg and 10mg/Kg Xylazine respectively), and transcardially perfused respectively with PBS and ice-cold 4% paraformaldehyde in PBS. Brains were removed and subsequently post-fixed overnight at 4°C in the same fixative solution. Coronal forebrain and rostral mid-brain slices were obtained by using a 7000SMZ-2 VIBROTOME (Campden Instruments) and collected as 50 μm floating sections. Slices were blocked for 90 min at room temperature (RT) in blocking solution made of PBS containing 0.3% Triton X-100 (SIGMA) and 5% Normal Goat Serum (NGS, Vector laboratories). Sections were incubated overnight (O/N) at 4°C with primary antibodies in PBS containing 0.3% Triton X-100 and 1% NGS in PBS. The following antibodies were used for immunofluorescence of fixed brains: human monoclonal anti-LGI1 (LRR1, LRR2 and LRR3) [6] at 12.8 µg/mL, 7.5 µg/mL and 6.7 µg/mL respectively, rabbit anti-AnkG (1:500), guinea pig anti-AnkG (1:500), mouse anti-synaptophysin, mouse anti-GFAP (1:400), mouse anti-PSD-93 (1:500). Afterwards, sections were washed three times (15 min each) in PBS containing 0.3% Triton X-100 and then incubated with the appropriate secondary antibodies (1:200) (2h, RT) in PBS containing 0.3% Triton X-100 and 1% NGS. Finally, sections were washed three times in a solution containing 0.3% Triton X-100 in PBS, incubated with DAPI (1.5 µg/mL; Sigma-Aldrich) for 10 min, washed one last time in PBS and mounted in Vectashield (Vector laboratories). Slides were kept at 4°C until use. Images were acquired on a Zeiss LSM-780 Confocal scanning microscope. All experiments involving WT and *lgi1^-/-^* comparisons were performed in parallel and the same acquisition settings were applied to both genotypes.

#### Hippocampal slice cultures and transfection of hippocampal neurons

Young Wistar rats (P7-P8) or mice (P7-P10) of either sex, were decapitated, the hippocampi were removed, and individual slices were collected (350µm) and placed on Millicell membranes (Merck-Millipore) inserted into 35mm Petri dishes containing 1mL of culture medium. The culture medium contained (in mL) 25 MEM, 12.5 HBSS, 12.5 horse serum, 0.5 penicillin (10,000U/mL), 0.5 streptomycin (10mg/mL), 0.8 glucose (1M), 0.1 ascorbic acid (1mg/mL), 0.4 HEPES (1M, pH 7.4), 0.5 B27, and 8.95 milli-Q H2O. For rat organotypic slices, 5 µM Ara-C was added to the slices starting at 1DIV to limit glial proliferation and removed at 3 DIV. For Gene Gun (Helios Gene Gun, Bio-Rad) transfection, Gene Gun bullets were prepared following the manufacturer’s instructions using 1-μm gold micro-carriers and a gold (mg) / cDNA (μg) ratio of 6/25. DIV 2-3 cultures were shot (1 μg DNA per bullet) at 120 psi. Single cell electroporation of CA3 pyramidal cells was performed at DIV 4-5 with an Axoporator 800A (Molecular Devices). Borosilicate pipettes (7-10 MΩ) were filled with the following solution (in mM): K-gluconate 120, KCl 20, HEPES 10, EGTA 0.5, MgCl2 2, Na2ATP 2, and NaGTP 0.3 (pH 7.4). Plasmids were diluted to a final concentration of 33 ng/µl in the pipette solution. Slices were placed in an extracellular solution containing (in mM): 125 NaCl, 26 NaHCO3, 3 CaCl2, 2.5 KCl, 2 MgCl2, 0.8 NaH2PO4, and 10 D-glucose, continuously bubbled with 95% O2–5% CO2. A loose-seal (25-40 MΩ) was formed between the pipette and the selected cell and then a train of -12 V pulses (pulse width: 0.5 ms, frequency: 50 Hz) was delivered for 500ms. When indicated, doxycycline (300 ng/mL) was added to the culture medium to induce protein expression under the control of Tet-on regulatory elements. The use of single cell electroporation technique in a *Lgi1^-/-^* background presents three main advantages, i) it limits the signal of a non-engineered version of LGI1 to a single neuron in a native environment, ii) it further validates the specificity of the monoclonal antibody (LRR1) iii) it provides an opportunity to co-express a soluble GFP to trace neuronal morphology and to follow the distribution of LGI1 clusters throughout the transfected neuronal processes.

#### Immunofluorescence of organotypic slices

Organotypic slices were rinsed in PBS at 37°C and immediately immersed in ice-cold 4% paraformaldehyde in PBS. Slices were fixed at 4°C for 15 minutes and subsequently incubated in PBS containing 50 mM NH4Cl (15 min, RT) in order to quench the remaining free aldehyde groups. Afterwards, the Millicell membrane containing the slices was cut with a scalpel, and the slices were blocked (O/N, 4°C) in PBS containing 0.5% Triton X-100 and 10% NGS. Then, slices were incubated with the primary antibodies (24h, 4°C) in a solution containing 0.5% Triton X-100 and 2% NGS in PBS. The following antibodies were used in these experiments: human monoclonal anti-LGI1 (LRR1, 12.8 µg/mL), rabbit anti-DENDRA2 (1:200), rabbit anti-GFP (1:500), guinea pig anti-AnkG (1:500), mouse anti-PSD93 (1:500). Afterwards, sections were washed three times (20min each) in PBS containing 0.5% Triton X-100 and then incubated with the appropriate secondary antibodies (1:200) (2h, RT) in PBS containing 0.3% Triton X-100 and 1% NGS. Finally, sections were washed three times (20min each) in a PBS solution containing 0.5% Triton X-100, incubated with DAPI (1.5 µg/mL; Sigma-Aldrich) for 10 min, washed one last time in PBS and mounted in Vectashield (Vector laboratories). Slides were kept at 4°C until use.

#### Immunofluorescence on HEK cells

Transfection of HEK cells and characterization of EPTP1 and 2 specificity were performed as described in [6].

#### Image analysis

For the quantification of the expression levels of LGI1 in forebrain and hippocampal regions, maximal projections of z-stacks consisting of 3 optical slices acquired with a 10x objective were used. Several ROIs of the same size were drawn in different brain regions and the average grey value immunoreactivities were normalized to the overall hippocampal signal (for different brain region comparisons), to dorsal CA3 (for the analysis of the dorso-ventral pattern of expression), or to CA3 Stratum Radiatum (for comparisons within the dorsal hippocampus). The mentioned regions were chosen for normalization as they showed the highest immunoreactivity for LGI1 for each type of comparison. For the analysis of the density of LGI1 clusters, 63x images were acquired as z-stacks and the LGI1 signal was used to detect 3D objects with the 3D objects counter plugin [1]. Clusters’ density was expressed as the number of LGI1 clusters/µm^3^. For the analysis of LGI1 expression at the AIS, several ImageJ plugins were used in combination in a custom-written ImageJ macro. Briefly, z-stacks of 10 to 20 optical slices were acquired with a 63x objective in the different hippocampal subregions (CA1, CA3 and DG), and AnkG signal was used to detect 3D objects with the 3D objects counter plugin [1]. The objects were then added to the 3D ROI manager [5] and a 3D binary mask was created. The ROIs in this binary mask were processed with binary operators (Dilate and Fill holes) and the selection mask was over-imposed to the LGI1 z-stacks. The LGI1 signal ascribable to the AIS was quantified as average grey values with the 3D ROI manager, and compared between different hippocampal regions. For visual clarity in Fig. 3, LGI1 signal not corresponding to the AISs was erased using the inverse of selection for the previously described 3D mask. Then, maximal projections of 10 optical slices z-stacks were rendered and presented along with the orthogonal views of each maximal projection. VolumeJ was used to render volumetric reconstructions. For colocalization analysis, z-stacks of 10 optical slices were background subtracted and analyzed using the Intensity Correlation Analysis (ICA) plugin [3]. Colocalization values were expressed as Manders’ overlapping coefficients (MOC) since the Ch1:Ch2 pixel ratio was ∼1 [4].

#### Mass spectrometry analysis and data processing

Immunoprecipitated proteins were concentrated on a stacking gel that was cut into small pieces, rinsed three times with 50% acetonitrile in water, reduced with DTT, alkylated with iodoacetamide and submitted for tryptic digestion overnight at 37°C in 25 mM ammonium bicarbonate. Peptides were extracted three times with a mixture of 50% acetonitrile and 0.1% formic acid then dried using a speedvac.

After resuspension with 2% acetonitrile and 0.1% formic acid mixture, samples were analyzed by mass spectrometry using a hybrid Q-Orbitrap mass spectrometer (Q-Exactive, Thermo Fisher Scientific, United States) coupled to a nanoliquid chromatography (LC) Dionex RSLC Ultimate 3000 system (Thermo Fisher Scientific, United States). Samples were trapped with a C18 PepMap 300 trap column (300 μm × 5 mm, C18, 5 μm, 300 Å) and desalted with 0.1% formic acid in water for 3 min at a flow rate of 20 μl/min. Peptide separation was performed on an Acclaim PepMap RSLC capillary column (75 μm × 15 cm, nanoViper C18, 2 μm, 100 Å) at a flow rate of 300 nl/min. The analytical gradient was run with various percentages of acetonitrile and 0.1% formic acid in the following manner: (1) 2.5–25% for 57 min, (2) 25–50% for 6 min, (3) 50-90% for 1 min, and (4) 90% for 10 min. MS spectra were acquired at a resolution of 35,000 within a mass range of 400–1,800 m/z. Ion accumulation was set at a maximum injection time of 100 ms. Fragmentation spectra of the 10 most abundant peaks (Top10 method) were acquired with high-energy collision dissociation (HCD) at a normalized collision energy of 27%.

All raw data files generated by MS were processed to generate mgf files and searched against the Swissprot database (Swissprot version june 2019, (560,118 sequence), with the taxonomy mus musculus, using the MASCOT software (www.matrixscience.com, version 2.3). Search parameters were as follows: mass tolerance: 10 ppm on parent ion and 0.02 Da on fragment ions, and a maximum of two missed tryptic cleavages. Methionine oxidation as well as carbamidomethylation of cysteine were considered as variable modification. The database was searched in the decoy mode. The emPAIs were automatically calculated by the MASCOT algorithm. All samples were analyzed in triplicates. For each antibody, interactomes from wild type samples were subtracted from the background interactomes obtained from LGI1 knockout samples using Excel and homemade python scripts. Partner proteins were considered positive when the emPAIs from WT samples were above 3 compared to those from *Lgi1^-/-^* tissue. Keratins, ribosomal proteins, transcription factors and immunoglobulins were excluded from the list of candidates. Candidate proteins were classified using the pantherdb web site (www.pantherdb.org).

---

**ESM_1.**
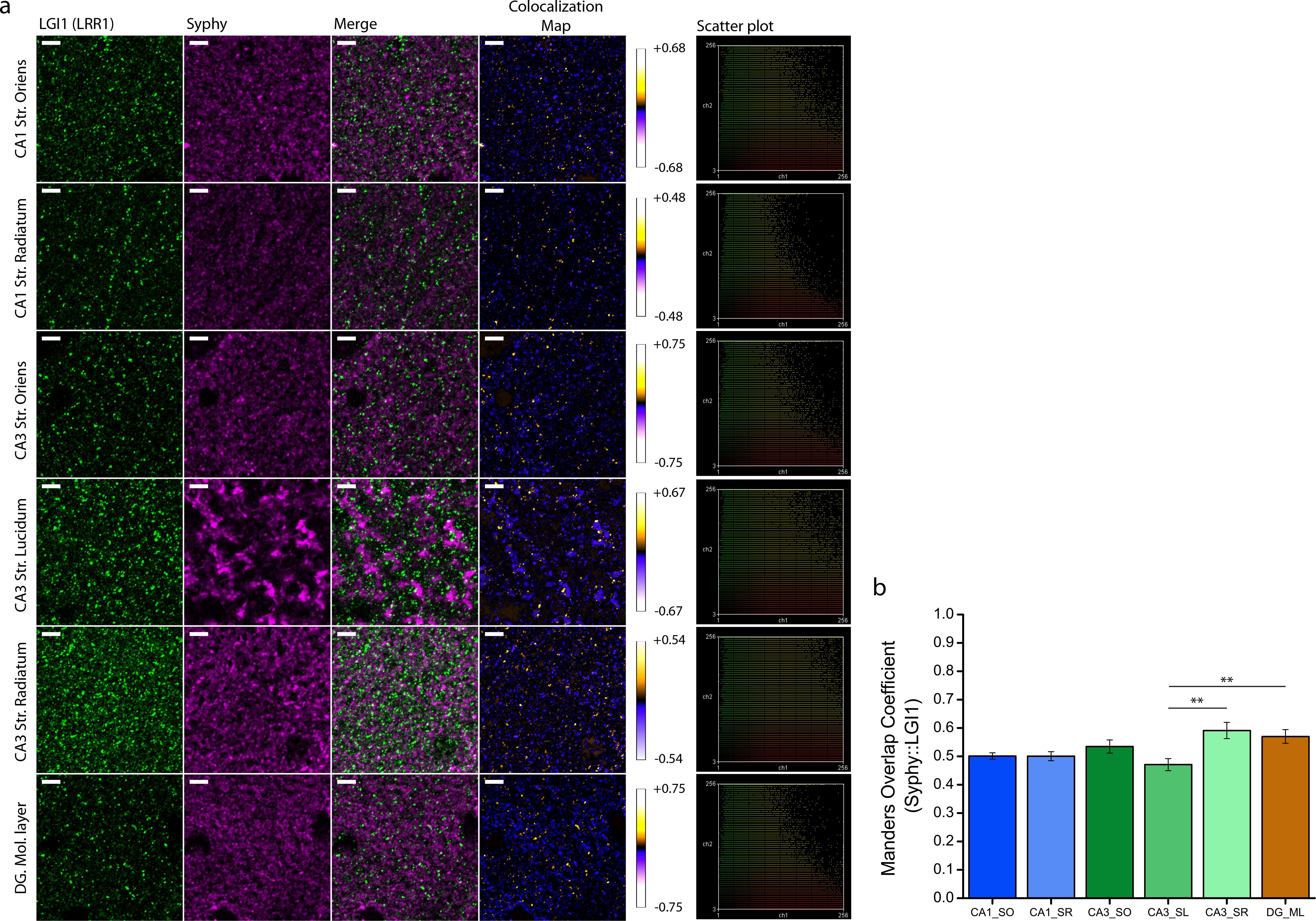
LGl1 partially colocalizes with the presynaptic marker synaptophysin in the hippocampus. a High­ magnification pictures of LGl1 (green, LRR1) and synaptophysin (magenta) in the different hippocampal strata. Colocalization maps are shown (right columns) with their respective colocalization index values. Scatter-plots illustrate partial colocalization in all the analysed regions. Scale bar = 5 µm. b Manders’ Overlapping coefficients for LGl1::synaptophysin colocalization in the different regions analyzed. Columns and error bars show mean ± SEM. n = 2 independent experiments (WT; LRR1 n = 1; LRR2 n = 1). For the sake of clarity only significant comparisons are shown. *\*p<0.05 ;* **p<0.01. Any other comparison is non-significant. n CA1 Str. Oriens = 8 Fields of view; ROls; 111 CA1Str. Radiatum = 8 Fields of view; n CA3 Str.Oriens = 8 Fields of view; n CA3 Str. Lucidum = 8 Fields of view; n CA3 Str. Radiatum = 8 Fields of view; n DG Molecular Layer = 8 Fields of view

**ESM_2.**
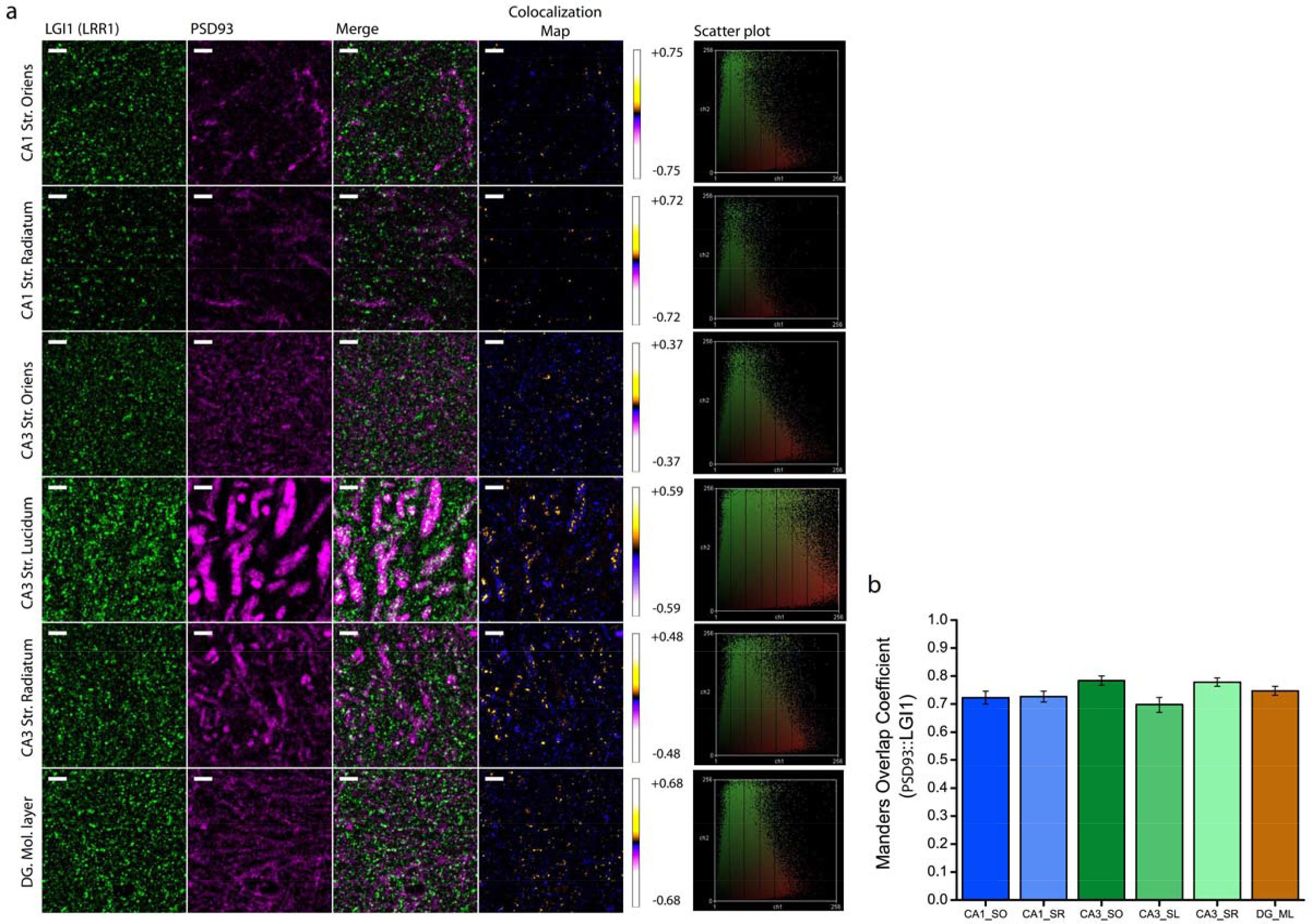
LG11 partially colocalizes with the postsynaptic marker PSD93 in the hippocampus. a High­ magnification pictures of LGl1 (green, LRR1) and PSD93 (magenta) in the different hippocampal strata. Colocalization maps are shown (right columns) with their respective colocalization index values. Scatter-plots illustrate partial colocalization in all the analysed regions. Scale bar = 5 µm. **b** Manders’ Overlapping coefficients for LGl1::PSD93 colocalization in the different regions analyzed. Columns and error bars show mean ± SEM. n = 2 independent experiments (WT; LRR1 n = 1; LRR2 n = 1). n CA1 Str. Oriens = 8 Fields of view; ROls; n CA1 Str. Radiatum = 8 Fields of view; n CA3 Str.Oriens=8 Fields of view; n CA3 Str. Lucidum = 8 Fields of view; n CA3 Str. Radiatum = 8 Fields of view; n DG Molecular Layer = 8 Fields of view

**ESM_3.**
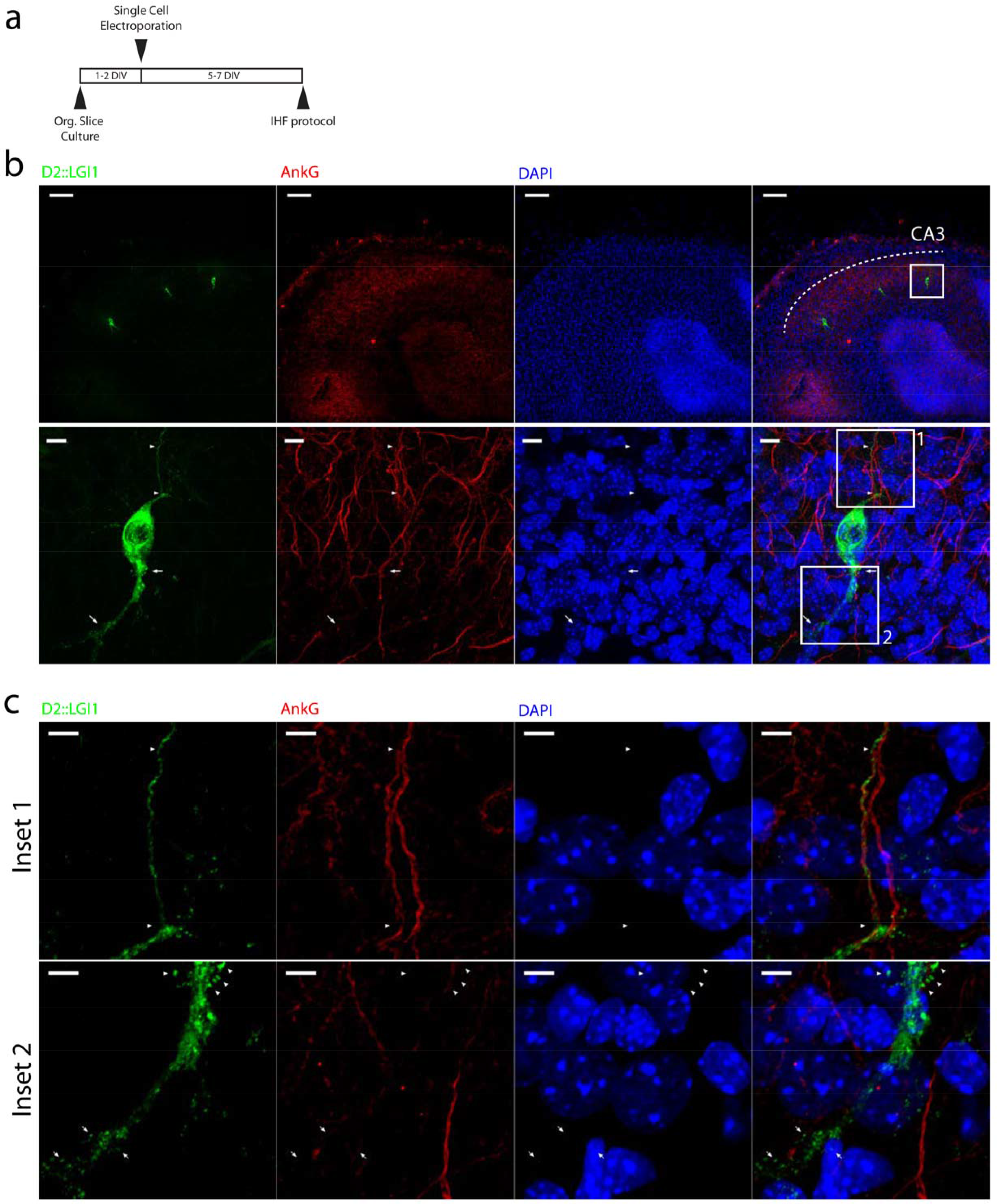
Subcellular distribution of DENDRA2::LGl1 studied by single cell electroporation and immunohistofluorescence in CA3 pyramidal neurons of mouse organotypic slices. a Schematic of the experimental protocol used for single cell electroporation and immunostaining. b Low-magnification picture of three indivridual CA3 pyramidal neurons electroporated with DENDRa2::LGl1 and immunostained with anti-DENDRA2 (green), anti-AnkG (red), and DAPI (blue). Scale bar = 100 µm (top panels). Detail of the inset outlined in the top panels showing DENDRA2::LGl1 expression in the AIS (arrowheads) and in the apical dendrites (arrows). Scale bar = 10 µm (bottom panels). c details of the insets 1 and 2 in (b). In the top panels, arrowheads point towards the AIS. In the bottom panels, arrowheads point towards thorny excrescences of the proximal apical dendrites. Arrows indicate the distal apical dendrite. Images are representative of 8 CA3 pyramidal neurons from 2 independent cultures.Scale bars = 5 µm

**ESM_4.**
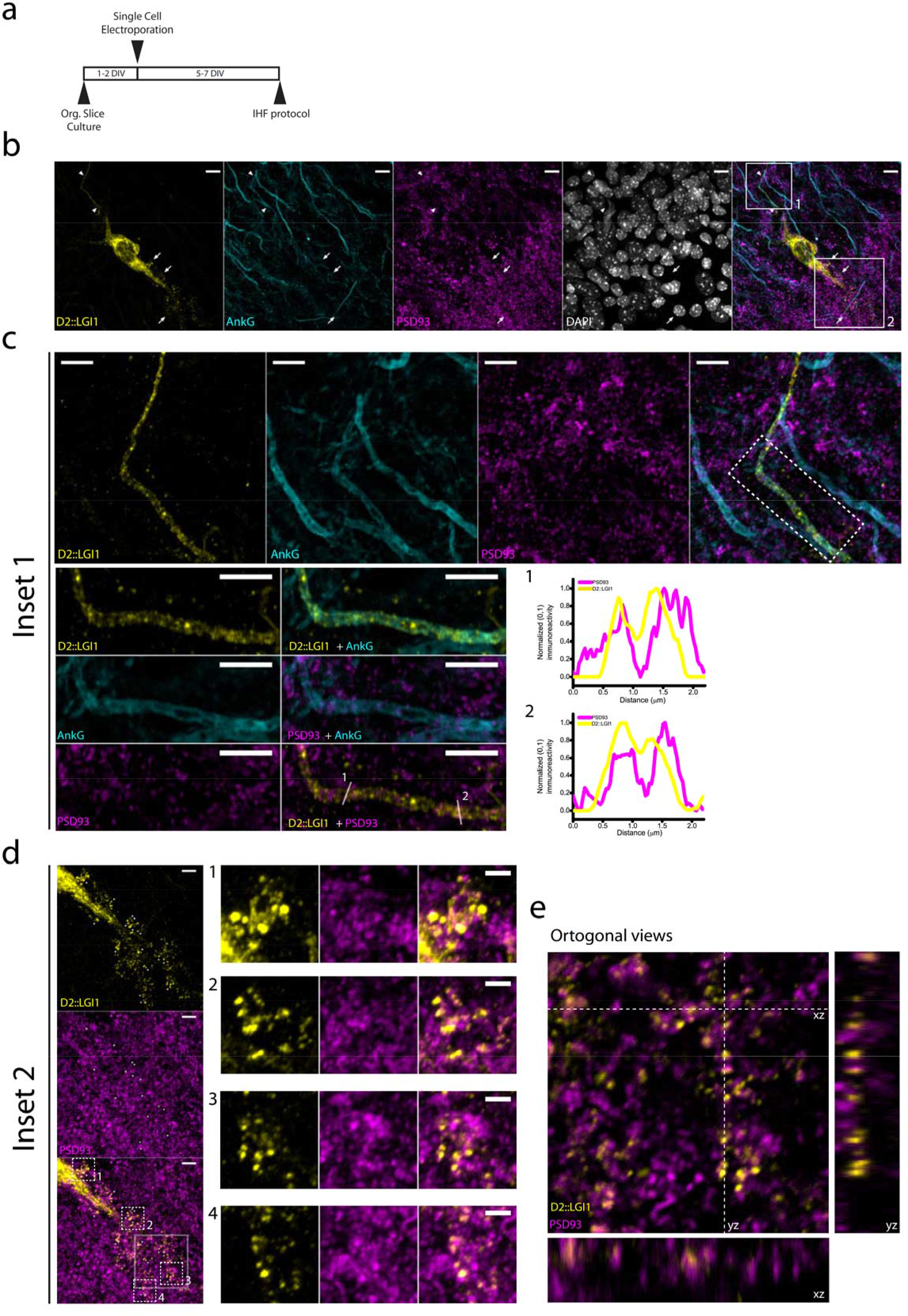
DENDRA2::LGI1 colocalizes with AnkG and PSD93 after single cell electroporation in mouse hippocampal slices. a Schematic of the experimental protocol used for single cell electroporation and immunostaining. b Low-magnification picture of one individual CA3 pyramidal neuron electroporated with DENDRA2::LGl1 and immunostained with anti-DENDRA2 (yellow), anti-AnkG (cyan), anti­ PSD93 (magenta), and DAPI (gray).Arrowheads indicate the Axonal Initial Segment. Arrows indicate thorny excrescences and the basal dendrite. Scale bar = 10 µm. c High magnification detail of the region (inset 1) outlined in b showing prominent expression of DENDRA2::LGl1 within the axonal initial segment (top panels), and colocalization of DENDRA::LGl1 clusters with PSD93 at the AIS. The boxed region in the top panels is shown in the bottom left panels. The line traces in the bottom right panels (1 and 2) correspond to the lines depicted in the left panel. Scale bar = 5 µm. d High magnification detail of the region (inset 2) outlined in b showing expression of DENDRA2::LGl1 at the thorny excrescences and the basaldendrite (left).Scale bar = 5 µm. Insets 1, 2, 3, and 4 (dashed lines) are depicted in the adjacent panels (right). Scale bar = 2 µm. e Orthogonal views of the region boxed (solid line) in d showing co-localization of DENDRA2::LGI1 with PSD93. Scale bar = 2 µm

**ESM_5.**
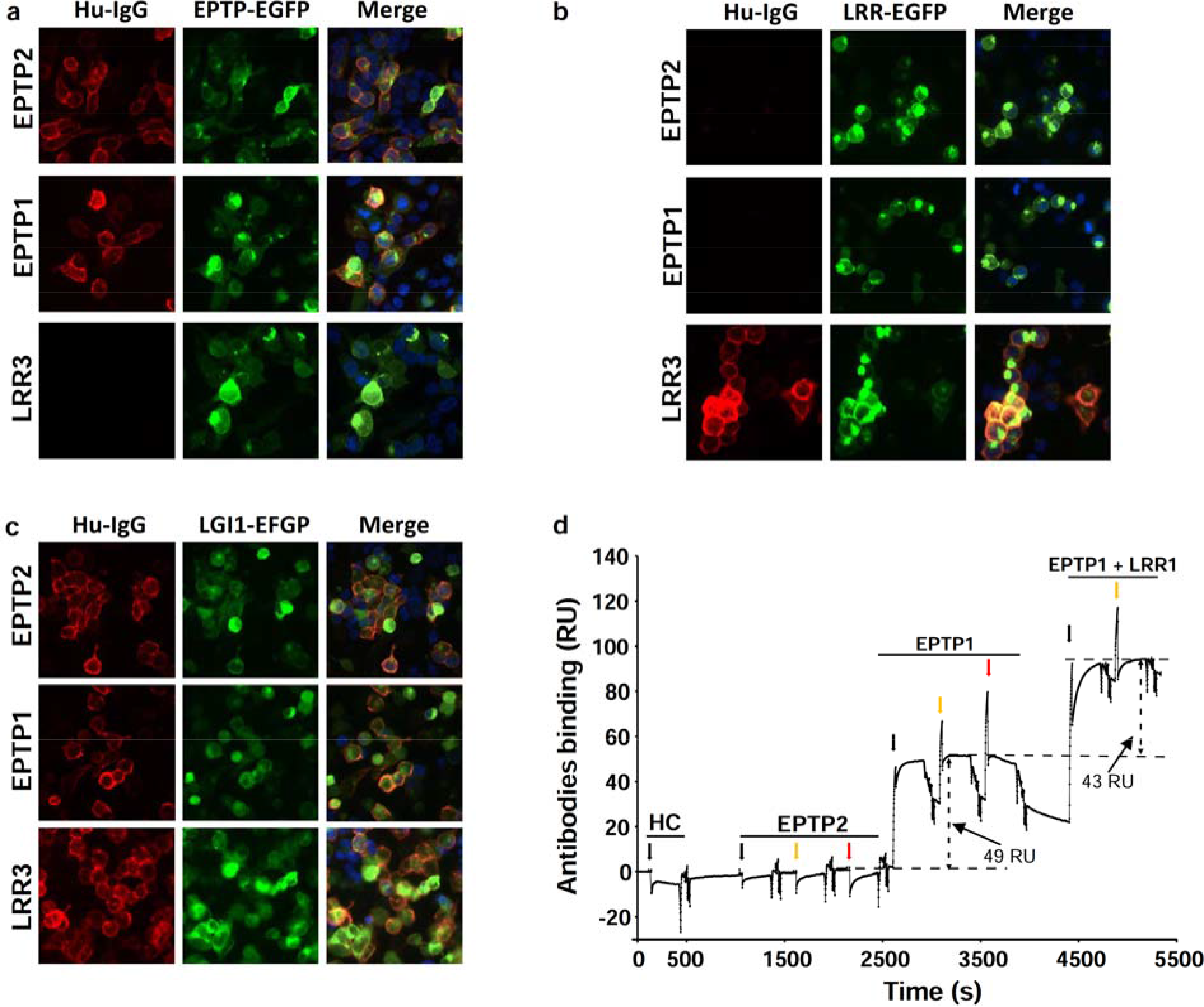
Characterisation of the anti-EPTP antibodies. Membrane tethered LGl1-EPTP (a) LGl1-LRR **(b)** or LGl1 (c) intracellularly fused to EGFP (EPTP-EGFP, LRR-EGFP, LGl1-EGFP respectively) were transfected into HEK293T cells and EPTP1, EPTP2 or LRR3 were applied to these live cells. Human lgG (Hu-lgG) binding is shown in red and EGFP fluorescence is shown in green. **d** 1-2 fmoles of LGl1 were captured on covalently immobilized EPTP2. Human control lgG (HC), EPTP2, EPTP1 and a mixture of EPTP1 and LRR1 were successively injected during 300s at 100 nM (black arrows), 200 nM (orange arrows) and 400 nM (red arrows). Horizontal dashed lines indicate the plateau level for each lgG. EPTP2 did not bind LGl1 indicating the absence of LGl1 multimeric forms. EPTP1 and LRR1 epitopes were similarly accessible leading to an increment of 49 and 43 RU respectively. Results are representative of 3 independent experiments.

**ESM_6.**
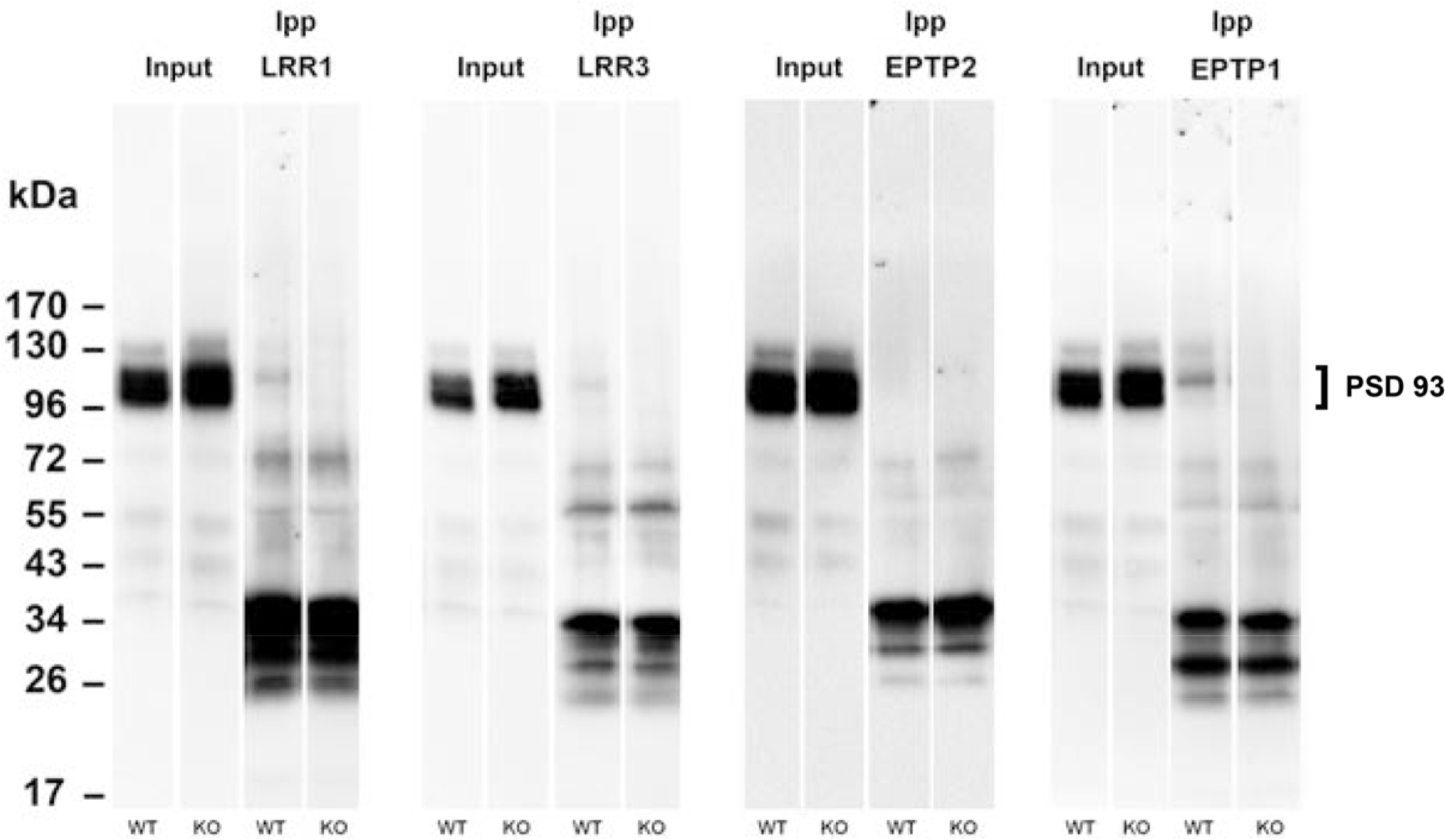
Specificity of PSD93 immunoprecipitation. Immunoprecipitations by LRR1, LRR3, EPTP2 and EPTP1 from wild type and *Lgi1^-/-^* samples were analysed by western blot using anti PSD93 antibodies. Bands between 26 and 34 kDa denote cross-reactivity of the secondary antibody with light chains of immunoprecipitating antibodies.

**ESM_’7.**
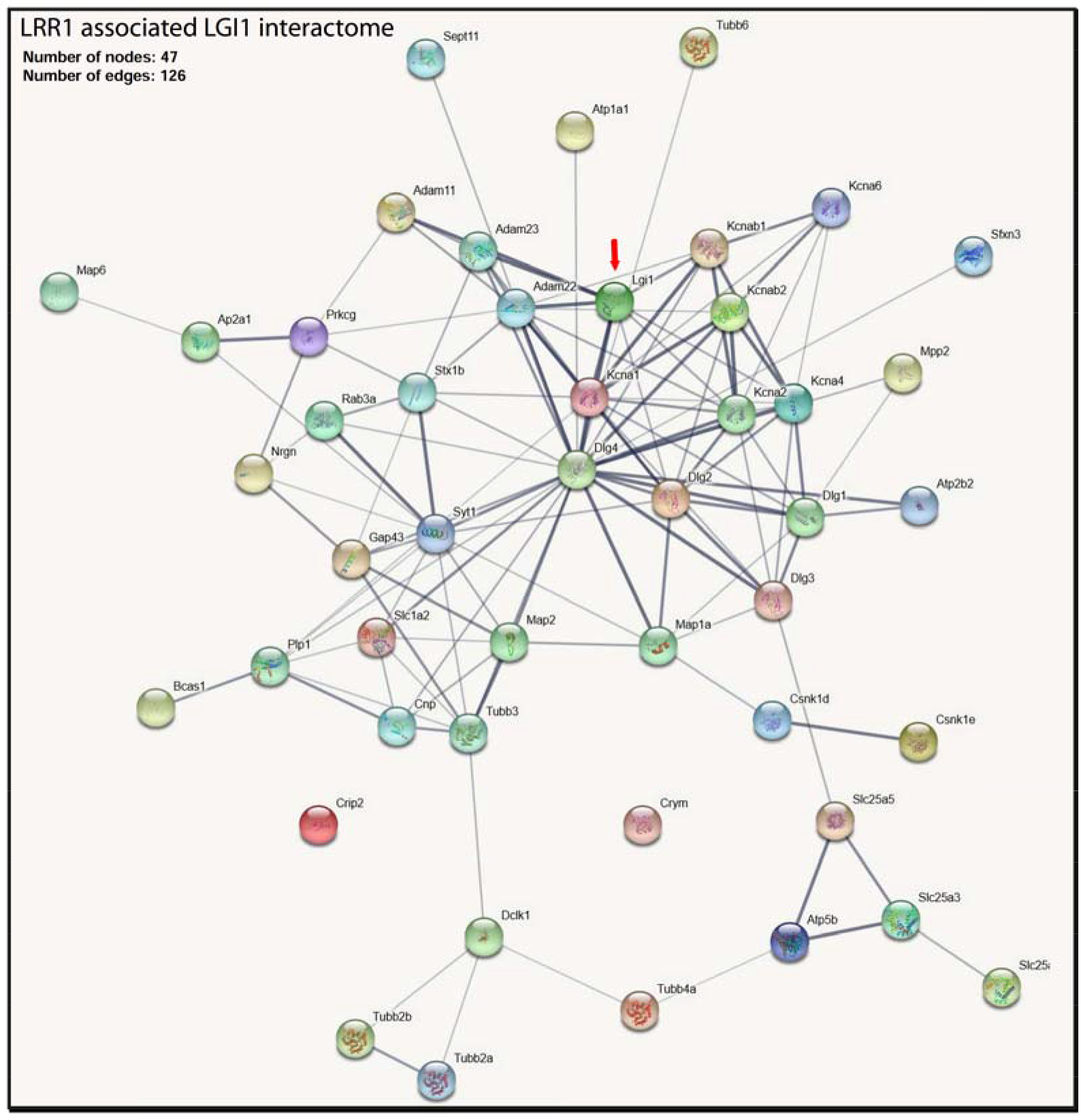
String Map of the LGl1 interactome associated with LRR1. Edge thicknesses indicate the strength of the data support

**ESM_8.**
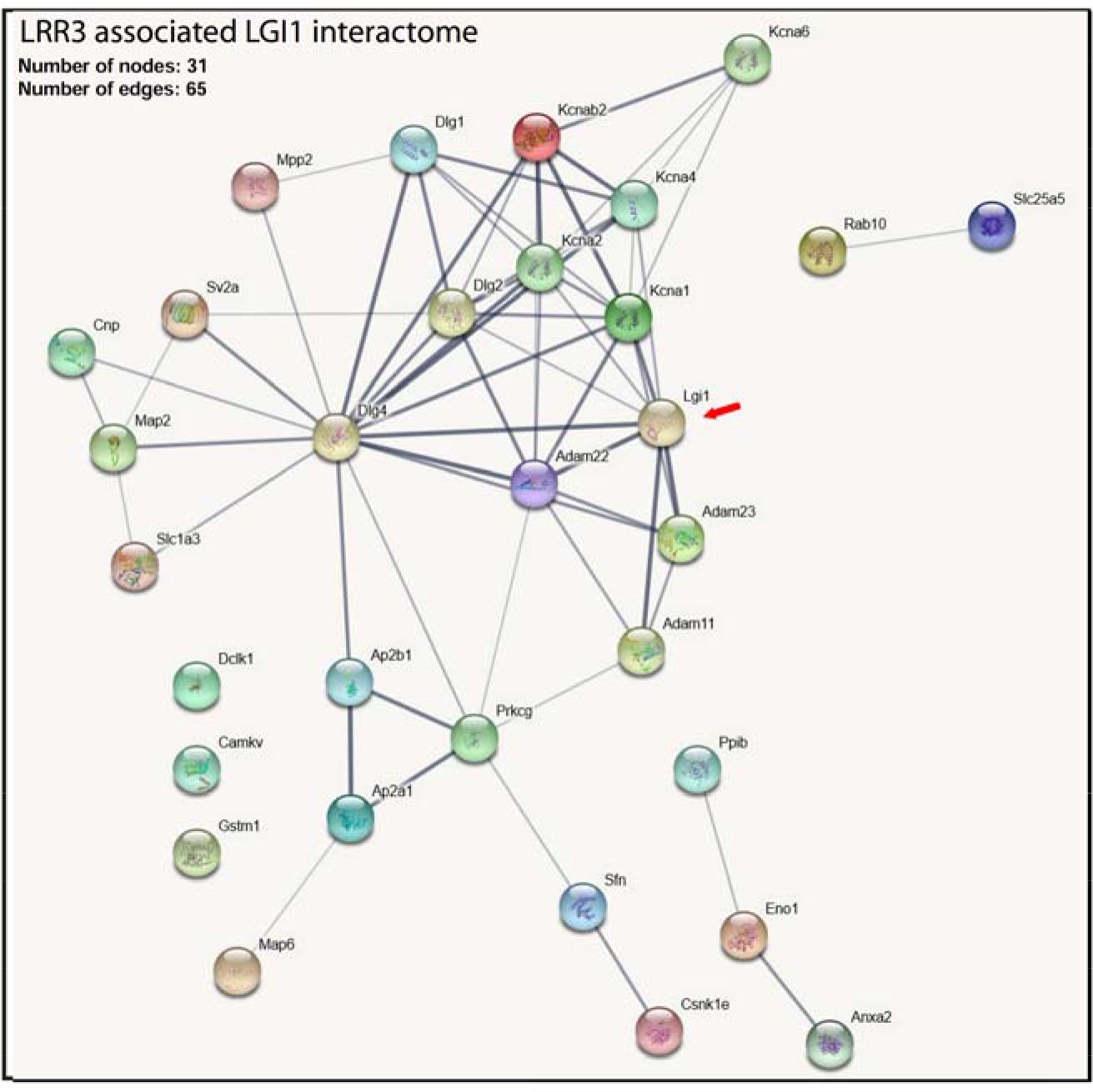
String Map of the LGl1 interactome associated with LRR3. Edge thicknesses indicate the strength of the data support

**ESM_9.**
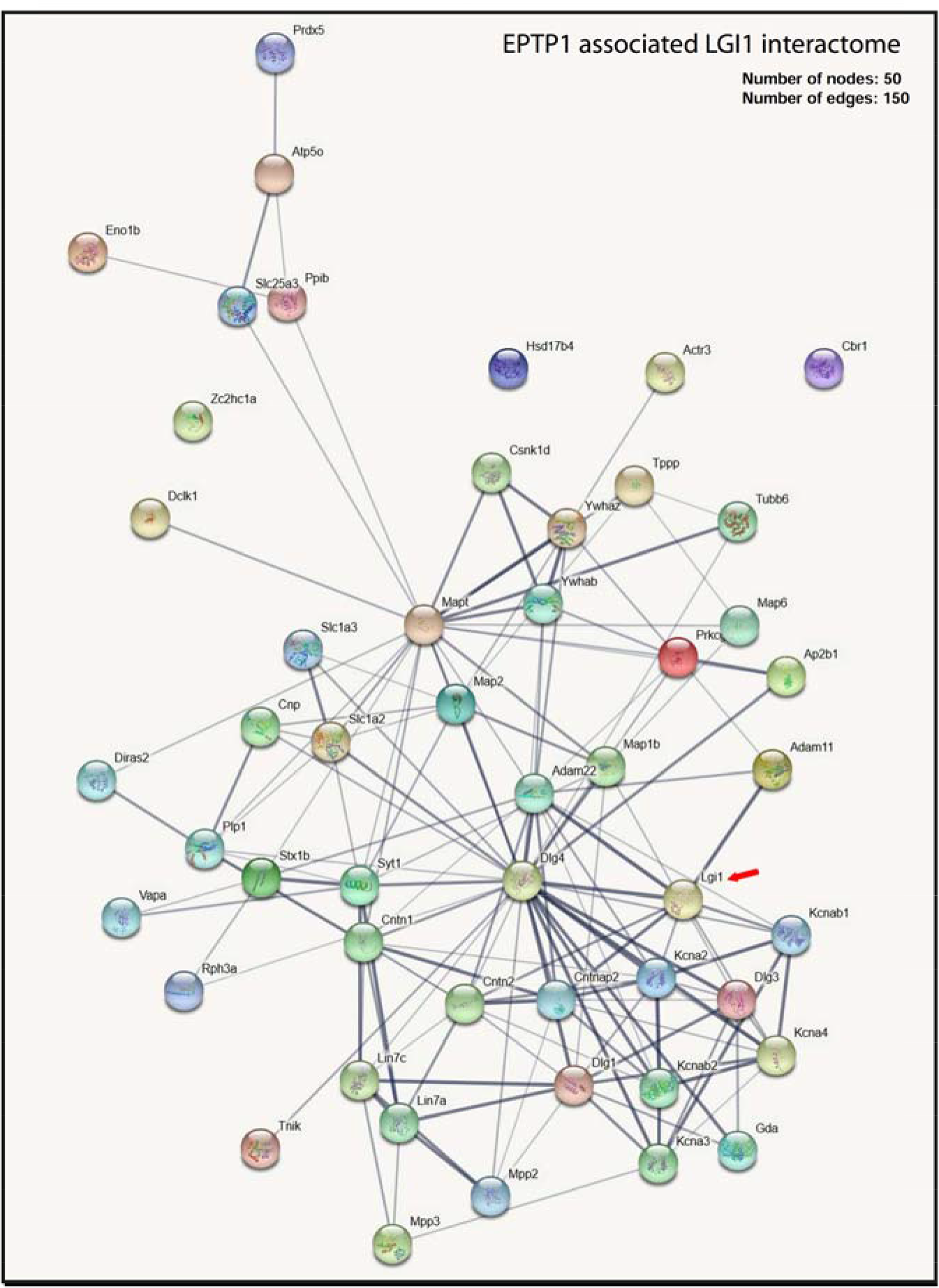
String Map of the LGl1 interactome associated with EPTP1. Edge thicknesses indicate the strength of the data support

**ESM_10.**
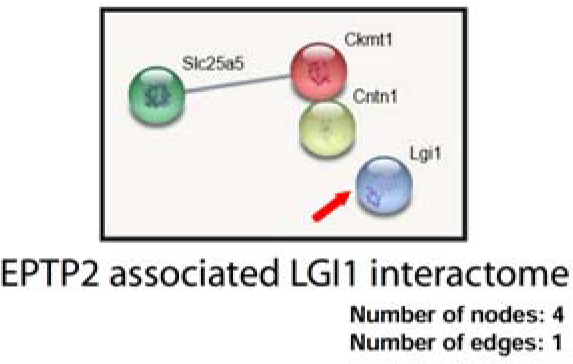
String Map of the LGl1 interactome associated with EPTP2. Edge thicknesses indicate the strengthof the data support

**ESM_11.**
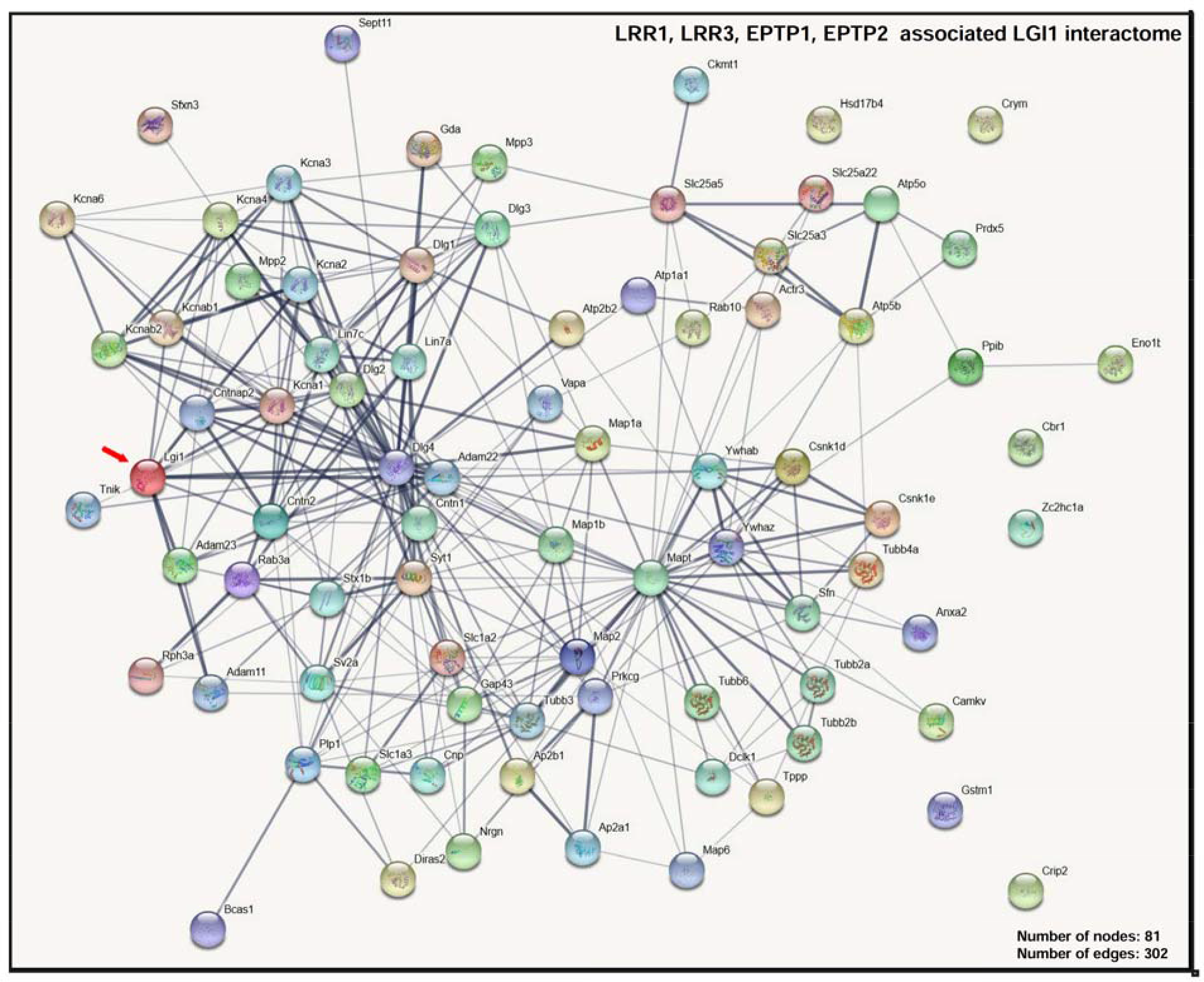
String Map of the LGl1 interactome associated with LRR1, LRR3, EPTP1 & EPTP2. Edge thicknesses indicate the strength of the data support

**Spreadsheet.**
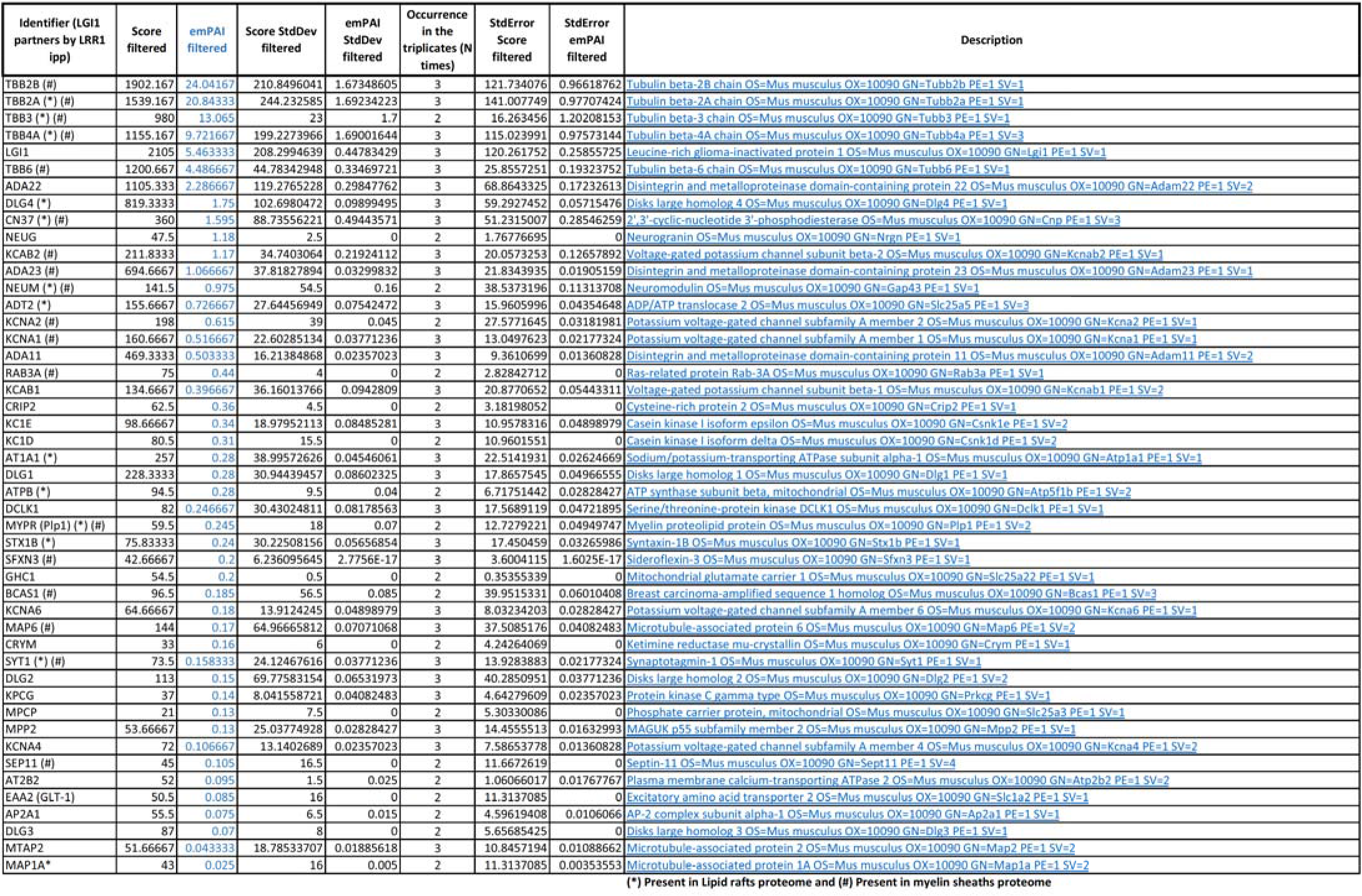
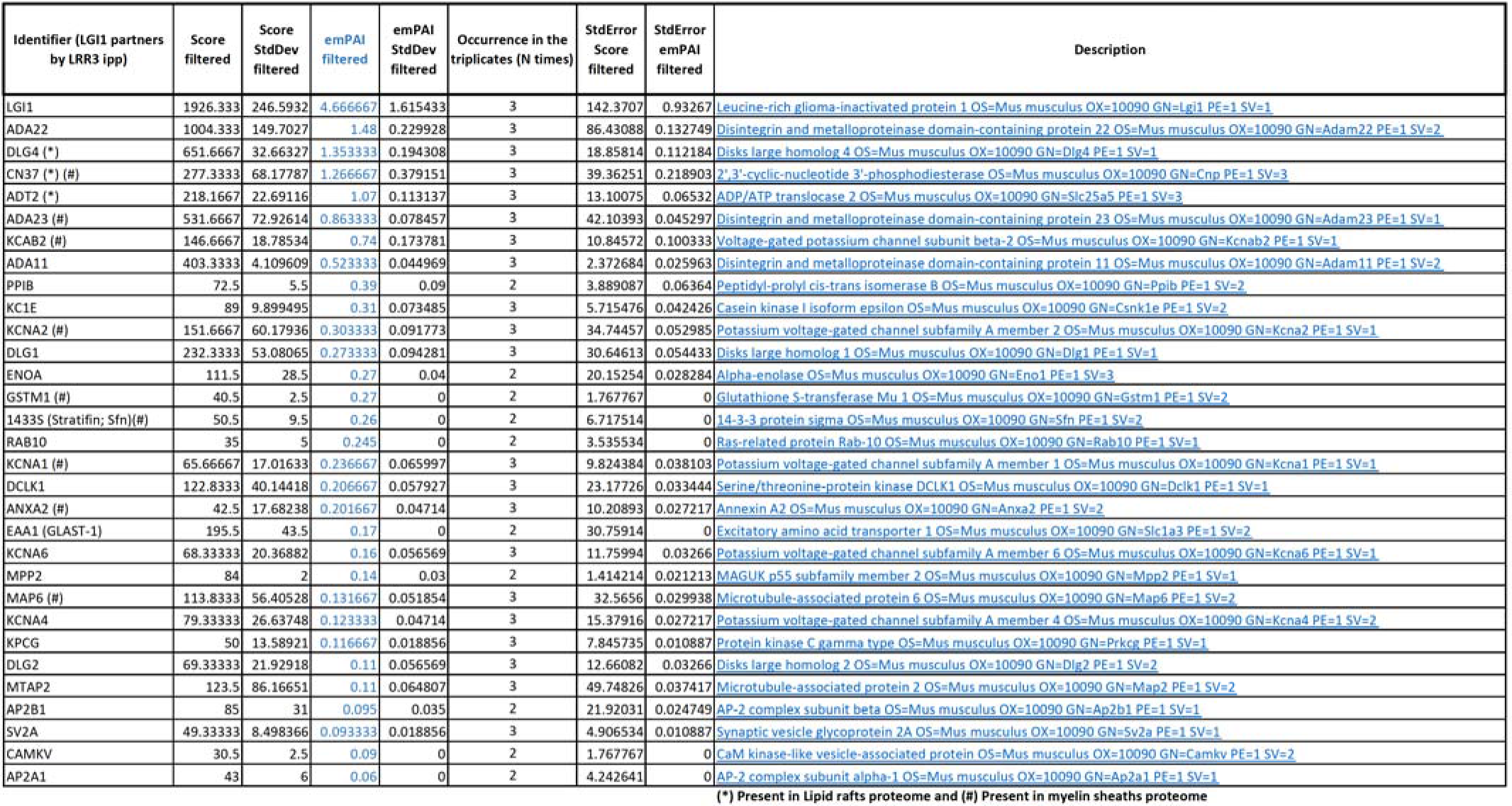
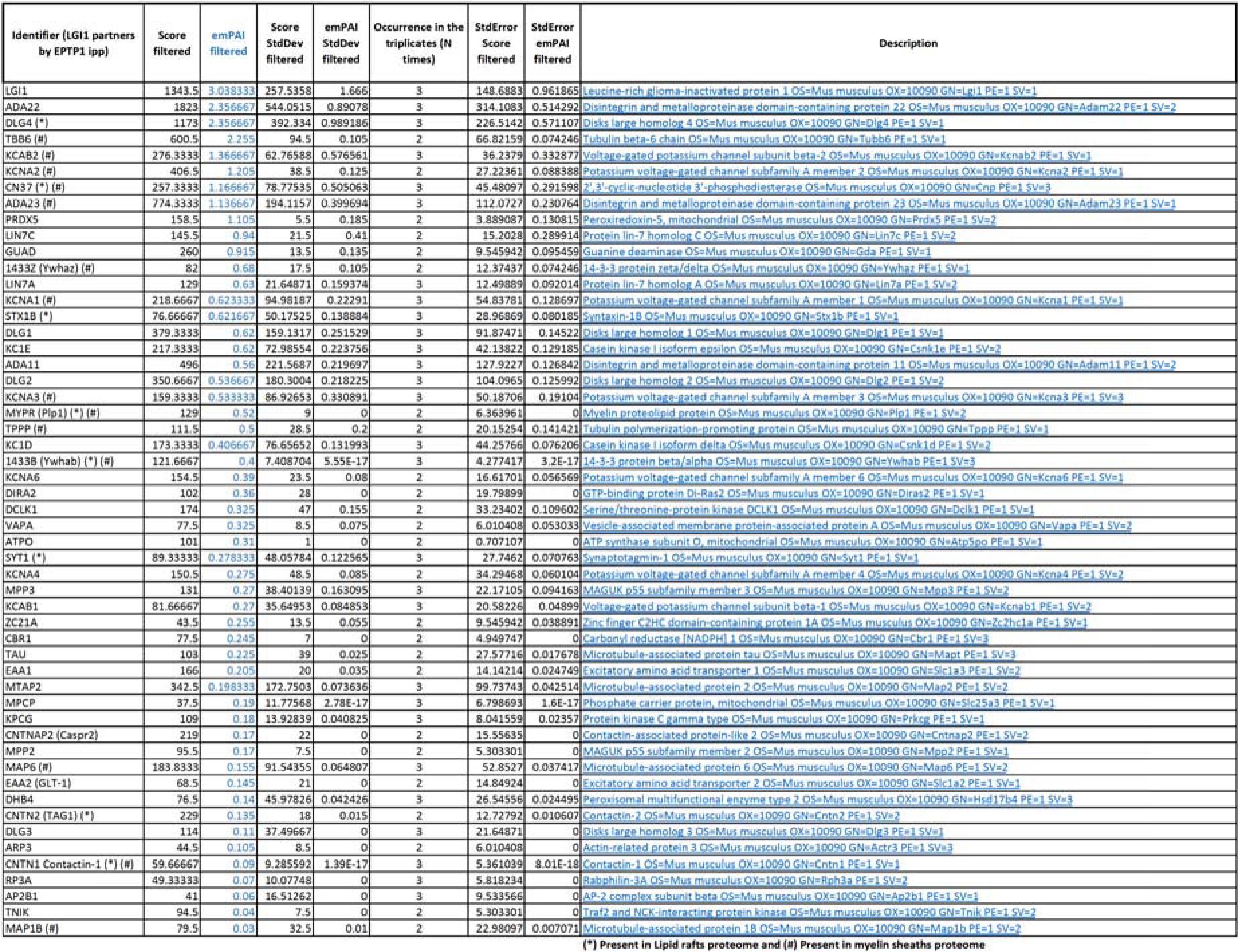
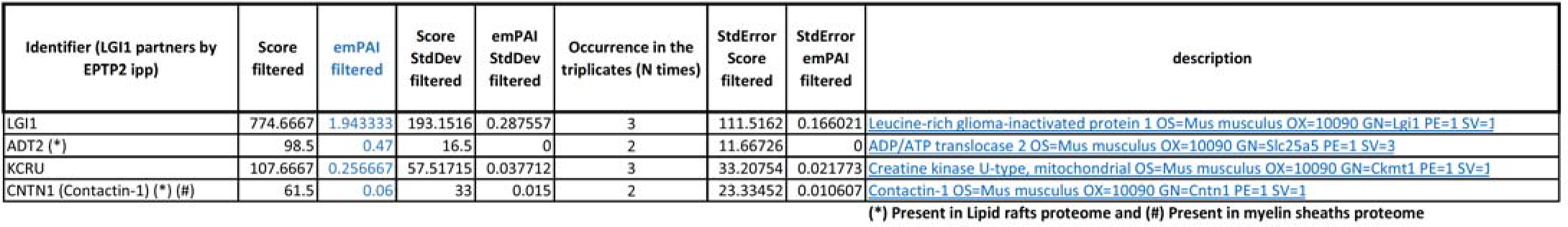
LGl1 partners immunoprecipitated by recombinant monoclonal antibodies derived from patients with LGI1-dependent limbic encephalitis. Partner proteins immunoprecipitated by each anti-LGl1are listed in the following 4 figures . Partners were considered as significantly present in experimental points if the emPAI ratio of the partner in the experimental point is superior to 3 compared to, when present, the emPAI in control *Lgi1-^1^-* controls. Scores and emPAls from control *Lgi1-^1^-* control points are already subtracted (filtered) from the reported scores and emPAls in the associated sheets. Keratin, immunoglobulins, ribosomal proteins and transcription factors were excluded from the analysis

## Notes

### Competing Interest Statement

The authors have declared no competing interest.

